# Selective brain network stimulation by frequency entrainment

**DOI:** 10.1101/2025.04.30.651283

**Authors:** Mónica Otero, Elida Poo, Felipe Torres, Cristobal Mendoza, Caroline Lea-Carnall, Alejandro Weinstein, Pamela Guevara, Jesus Cortes, Pavel Prado, Joana Cabral, Wael El-Deredy

## Abstract

**Background:** Brain stimulation at specific frequencies has been shown to have therapeutic effects for certain neurological disorders. However, it remains unclear how these oscillatory inputs interact with neuronal dynamics and what drives this frequency selectivity. Here, we propose that this is achieved by targeting specific brain circuits through frequency-selective network entrainment.

**Methods:** To demonstrate that this type of entrainment can occur in the brain connectome structure, we use a minimal physics-based model of coupled oscillators that preserves only the essential structural connectivity of real brain networks. Using this model, we test how periodic stimulation influences network oscillatory dynamics, identifying sub-networks that synchronize with specific stimulation frequencies. Furthermore, we validate the suitability of the model to reproduce well-known frequency-selective entrainment using visual and auditory periodic stimulation.

**Results:** Our model reproduces selective network entrainment, whereby distinct stimulation frequencies favor the synchrony of different sub-networks. Crucially, entrainment exhibits a physics-predicted inverse relationship with frequency: lower-frequency stimulation favors broader synchrony across the network, while higher-frequency inputs produce spatially confined effects. These patterns emerge purely from network structure and oscillatory physics, independent of biological details.

**Conclusion:** This physics-based approach elucidates the fundamental mechanistic principles governing frequency-selective brain network stimulation. Our minimal model demonstrates that selective entrainment of specific subnetworks can be achieved through frequency selection alone, largely determined by network structure and oscillatory dynamics. These findings provide a theoretical foundation for understanding how network architecture determines stimulation selectivity, supporting the development of principled approaches to targeted neuromodulation.

**Author summary:** In this work, we explore how external rhythmic stimulation interacts with the brain’s own rhythms. We use a physics-based model that represents the brain as a network of interconnected oscillators, linked according to the human connectome. Through this model, we show that stimulation effects are not limited to the targeted region but spread across the network, depending on the stimulation frequency, its spatial location and the underlying network dynamics. Our results reveal that low-frequency rhythms promote large-scale synchrony, while high-frequency rhythms engage more localized areas, all emerging from the brain’s structural connections and communication delays.

The simulated cortical activation exhibits a frequency-selective profile that aligns strongly with that estimated from sensory-evoked electroencephalographic activity. This agreement suggests that differences between individuals in how their brain networks operate may explain why stimulation affects each person differently. Our findings offer a mechanistic framework for designing personalized stimulation protocols, illustrating how computational modeling can guide the optimization of noninvasive brain stimulation in both research and clinical contexts.

## Introduction

Neural oscillations are thought to play a crucial role in coordinating neural activity, allowing the brain to integrate information across different regions and timescales [1]. This integration supports core cognitive processes such as perception, attention, and memory [2–4]. At the local circuit level, oscillatory activity is commonly observed in local field potentials (LFP), with representative frequencies around 40 Hz [5, 6].

Oscillatory activity at the macroscopic level is generally slower and strongly influenced by network interactions [7], contributing to complex brain dynamics [8, 9]. The interplay between local intrinsic dynamics and global connectivity structures results in a metastable system, where brain networks exhibit transient synchronization patterns that can rapidly switch between different states [9–11], supporting the brain’s ability to adapt to changing cognitive demands and environmental conditions [12, 13].

Neurological disorders often manifest as network dysfunction — aberrant connectivity and altered oscillatory synchronization — that can improve with targeted resynchronization. For example, neurodegenerative syndromes selectively affect intrinsic large-scale networks, consistent with network-level pathology [14, 15]; schizophrenia exhibits abnormal beta/gamma synchrony linked to cognitive symptoms [16–18] and after stroke, behavioral deficits are often better predicted by disrupted functional connectivity than by lesion location, underscoring network failure as a driver of impairment [19–21]. Epilepsy likewise exemplifies pathological synchrony propagating through cortical networks [22, 23]. Conversely, non-invasive, frequency-tuned stimulation that re-synchronizes long-range interactions (e.g., theta tACS) can acutely enhance cognition in older adults [24, 25], and both human and animal studies demonstrate genuine oscillatory entrainment by tACS [26, 27].

More generally, periodic stimulation techniques, such as transcranial alternating current stimulation (tACS), repetitive transcranial magnetic stimulation (rTMS), theta burst stimulation (TBS), and repetitive sensory stimulation (e.g., visual, auditory, or tactile), have been shown to modulate oscillatory brain activity [28–32]. It is thought that repetitive stimulation entrains neural networks, restoring or reconfiguring dysfunctional neural synchrony associated with neurological and psychiatric disorders [33–35]. For example, rTMS at gamma frequencies (30-80 Hz) has shown potential in reducing symptoms of schizophrenia [33], while tACS targeting alpha oscillations (8-12 Hz) has been used to alleviate symptoms in major depressive disorder [34]. Furthermore, sensory stimulation, such as rhythmic auditory stimulation, has been shown to improve motor function in Parkinson’s disease by entraining beta oscillations (13-30 Hz) [35].

However, the outcomes of neurostimulation are variable, and there is no consensus on stimulation protocols as the underlying mechanisms that govern entrainment and signal propagation within large-scale brain networks are not fully understood [36, 37]. Despite recent advances, the challenge remains to identify optimal stimulation parameters tailored to individual neurophysiological profiles. Gaining a deeper insight into how stimulation parameters, such as frequency and stimulation site, interact with ongoing oscillatory brain dynamics is crucial for optimizing possible therapeutic interventions. An effective strategy to address this challenge is the use of computational models, which enable the systematic exploration and refinement of the theoretical range of stimulation parameters essential to achieve effective neurostimulation [38].

Computational models of whole-brain networks, e.g. [39] [40] [41], that simulate the interplay between local oscillatory activity and large-scale connectivity, provide a controlled environment to investigate the mechanisms of network entrainment. Connectome-coupled node dynamics show that conduction delays and topology shape a characteristic power spectrum with discrete frequency peaks (reflecting oscillatory modes) in which coherent metastable subnetworks form and dissolve [9, 11, 42]. In these models, the nodes transition between different functional configurations, each associated with a distinct oscillatory mode, establishing a direct correspondence between network structure and oscillatory dynamics [11, 42]. Network size and connectivity are further determinants in setting the dominant cortical frequencies [43], providing principled targets for frequency-specific entrainment. Computational models thus provide a framework for understanding how stimulation modifies the oscillatory dynamics by favoring specific modes and their underlying network configuration.

In this study, we propose that periodic stimulation works by selectively entraining oscillatory modes embedded in the brain’s structural network, synchronizing specific subnetworks depending on stimulation frequency. Using a connectome-based model of coupled oscillators, we examine how external rhythmic forcing drives frequency-specific patterns of synchronization and propagation across the brain. We assess whether these modeled patterns align with empirical EEG recordings obtained during visual and auditory rhythmic stimulation, thereby validating the model’s ability to capture the fundamental mechanisms of frequency-selective entrainment.

## Materials and methods

We implemented a Forced Kuramoto model to simulate brain dynamics under external stimulation using a network topology derived from human diffusion tensor imaging (DTI) data. To achieve this, we modified the classic Kuramoto model [39] by incorporating time delays [44] and introducing an external periodic force [45] applied to specific individual nodes. We examined how stimulation site and frequency influenced entrainment patterns in brain networks are affected by structural network parameters, namely global coupling and mean delay.

We initially identified the model parameters that maximized metastability (determined as the point of maximal spectral entropy, as in [11]) by exploring the parameter space defined by global coupling strength and mean delay. We tested the effect of stimulation frequency and site by systematically driving each node individually, and identified coherent networks that emerged as a result of this entrainment.

### Brain network model with Kuramoto oscillators

To explore the spontaneous dynamics of coupled brain areas, we used a variant of the classic Kuramoto model [39] incorporating time delays between connected brain regions [44, 46]. The dynamical equation that describes the behavior of the phase *θ_n_*(*t*) of a node *n* at time *t*, is represented by:

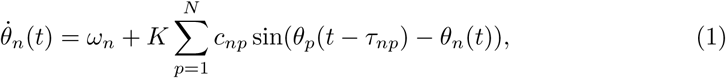

where *ω_n_* is the natural frequency of the oscillator *n*, assumed to be identical across all oscillators, and set to *ω_n_* = 40 Hz. The parameter *K* represents the global coupling strength between oscillators, *c_np_* denotes the coupling strength from node *p* to node *n*, *τ_np_* is the corresponding time delay. The use of 40 Hz as the intrinsic frequency for local oscillators is a modeling convention grounded in experimental neuroscience. It reflects the dominance of gamma oscillations commonly observed in cortical dynamics and provides a biologically plausible regime for investigating neural synchronization and emergent large-scale network behavior [47]. Subsequent computational studies [9, 11, 46] have maintained this assumption to simplify model design while effectively capturing the essential features of gamma-band neural synchrony. Additionally, using 40 Hz facilitates comparability across modeling studies and alignment with empirical MEG/EEG data.

In this study, we used the structural brain network from [46], constructed using the AAL90 parcellation template [48], which segments the brain into 90 regions that define the network oscillators or nodes. Connectivity was derived using probabilistic tractography, which estimates white matter tracts between brain regions [49] using DTI data of 21 healthy participants [46]. This connectome was initially used by [46] to model delayed interactions leading to frequency-specific amplitude modulations that resemble MEG resting-state networks. More recently, [11] employed the same structural network to study metastable dynamics and the spontaneous emergence of coherent subnetworks at distinct frequencies.

The network structure is represented by two *N* -by-*N* matrices: the connection weight matrix (*C*), where each element is proportional to the number of white matter tracts detected between two regions, and the inter-regional delay matrix (*D*), which accounts for transmission latencies between brain regions. The elements *c_np_* and *d_np_* represent the connection weight and delay, respectively, between two brain regions *n* and *p*.

To better understand the role of delays in the system, a mean delay factor (MD) is introduced, corresponding to the mean of the delay distribution. Since the distances between nodes remain fixed, varying MD is equivalent to modifying the conduction velocity (*v*) of neuronal signals [46], allowing the delays to be expressed as:

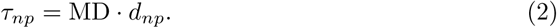

To characterize the system’s dynamics, we quantify metastability using spectral entropy [11, 50, 51]. A spectrum with more oscillatory modes will have higher spectral entropy than one with fewer modes [52]. This property is observed in EEG studies, where deep sleep or seizures show lower spectral entropy, than wakefulness [11, 50, 51].

Node spectral entropy (SE) is calculated using Shannon’s entropy formula, where power values are normalized and treated as probabilities P as a function of frequency bin *f_i_*:

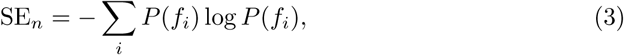

System spectral entropy is summed across all nodes:

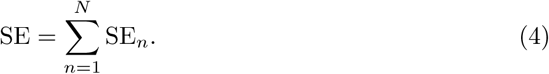

Simulations were conducted for 300 seconds using Eq. 1 to identify the set of parameters that maximize spectral entropy, SE (broader parameter space fully mapped in S1 Fig, as reported by [11]). In this point, highlighted by the yellow square (*K* = 4, MD = 21 ms) in Fig 1, the system exhibits the largest number of oscillatory modes, each peaking at a distinct frequency, namely *f* = 13.0, 15.0, 29.4, 41.2, 43.0 Hz. The lower panel of Fig 1 represents the power spectral density at this maximum entropy point. These frequency peaks are associated with limit-cycle attractors, meaning that the synchronised networks at these frequencies tends to persist for extended periods, reflecting the multi-state metastable dynamics of the network [11].

**Fig 1.**
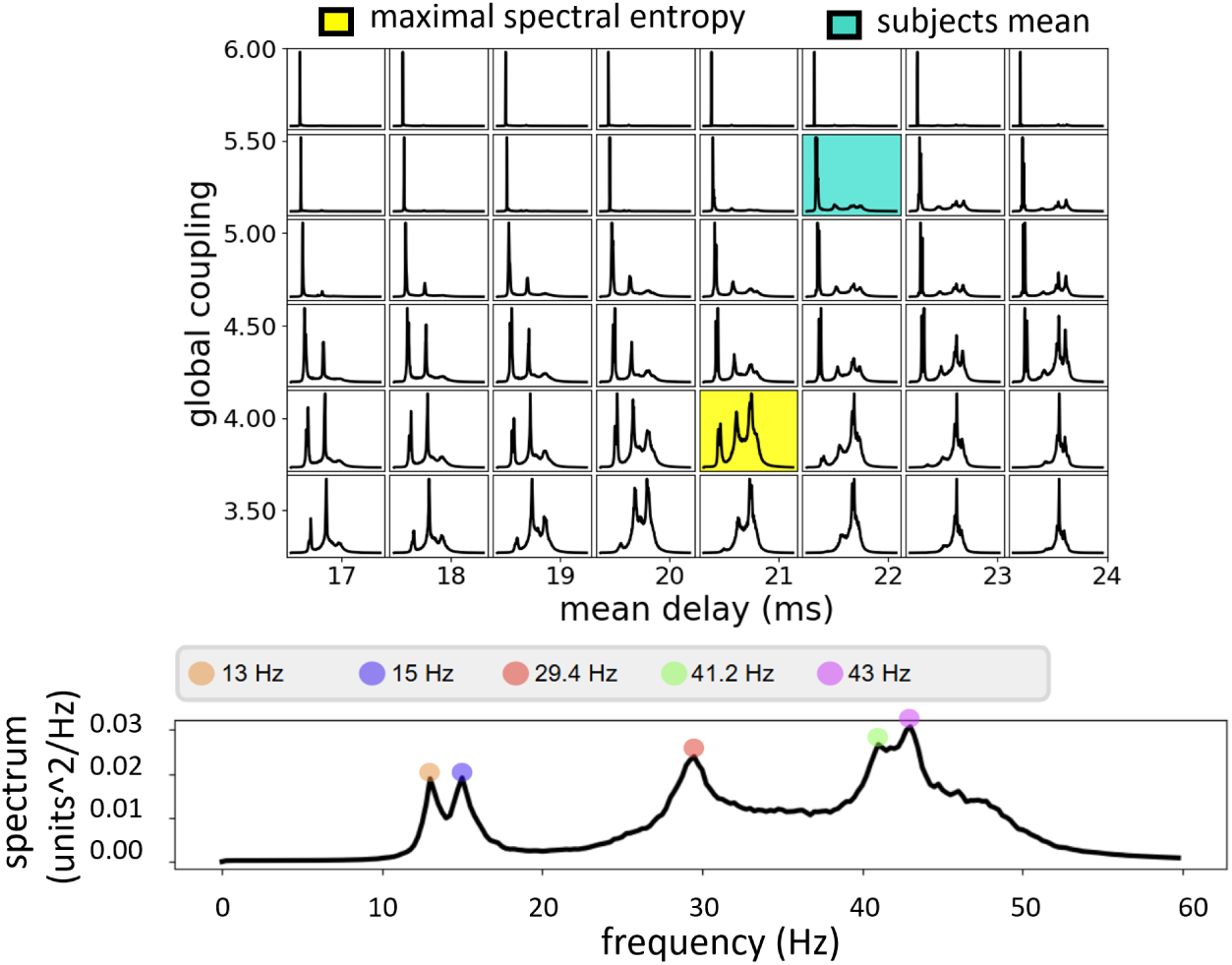
Multistate metastability in Kuramoto whole-brain network model. Top: Average spectra as a function of *K* and MD. The yellow square indicates the parameter set OP maxSE *K*, MD = 4, 21 ms corresponding to maximum spectral entropy. Cyan square indicate the model’s parameter space whose spectral entropy matched the mean spectral entropy of real EEG data from 32 healthy participants: OP EEG *K*, MD = 5.5, 22 ms. Bottom: Average spectrum at maximum spectral entropy. The peaks agree with the spontaneous oscillatory modes that emerge from the model dynamics with parameter set to 4, 21ms, as reported by Torres et al. 2024, which in turn map onto specific frequency-dependent functional networks ( [9, 11, 46]).

### EEG data processing and mapping to Kuramoto spectral space

To illustrate how sensitivity to stimulation depends on baseline network properties, we use eyes-open EEG recordings. Specifically, we examined how baseline spectral characteristics influence the response to stimulation by fitting the model spectrum to those obtained from experimental EEG recordings.

To do so, we analyzed two EEG datasets that shared the same resting-state acquisition and processing. EEG was recorded in thirty-two participants (17 males, 15 females; 26 ± 3 years), with normal or corrected-to-normal vision and no history of neurological/psychiatric disorders, participated under ethics approval from the Universidad de Valparaíso (CEC170-18 and CEC230-21) with written informed consent. EEG was acquired using a 64-channel BioSemi system at 8 kHz and preprocessed (average reference; 0.1–200 Hz 8th-order zero-phase Butterworth band-pass filter; downsampled to 512 Hz; ocular/motion artifact removal via ICA), then segmented into 5 s epochs.

To link baseline spectra to model responses, we compared each participant’s mean power spectral density (PSD; Welch method with 5 s Hann windows, 50% overlap; 0.2 Hz resolution) averaged across electrodes with mean model PSDs averaged across nodes across a grid of parameters {*K*, MD}, following [11]. Similarity between empirical and model spectra was quantified using the Jensen–Shannon distance (JSD; 0 = identical, 1 = maximal divergence) [53].

Mathematically, the Jensen-Shannon distance between two probability distributions *P* and *Q* is given by:

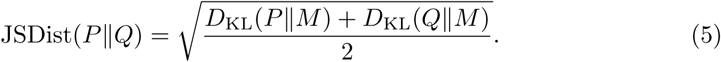

where *M* is the pointwise mean of *P* and *Q*, and *D*_KL_ represents the Kullback–Leibler divergence between two distributions (for example, between *P* and *M* , or *Q* and *M* ), defined as:

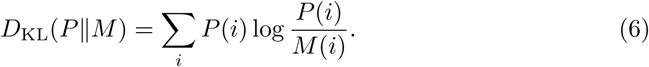

Using this approach, we identified parameter combinations (operating points of the model) that closely matched the model and participants’ EEG spectra, which can be interpreted as individual fingerprints reflecting subject-specific oscillatory dynamics. We hereafter refer to the EEG-derived operating points as OP EEG. The mapping of the model spectra to real EEG data is shown in S2 Fig and S3 Fig. The operating points OP EEG of different participants were: {*K*, MD} = {5, 19 ms}, {5, 20 ms}, {5.5, 22 ms}, {4.5, 18 ms}, {4.5, 19 ms} and {5, 21 ms}) . The operating point {*K*, MD}={5.5, 22 ms} also corresponds to the mean of the participants’ OP EEG parameters (cyan square in Fig 1).

### Modelling stimulation using Forced Kuramoto

To model stimulation we include an external periodic force that targets a single node based on [45]. The governing equation becomes:

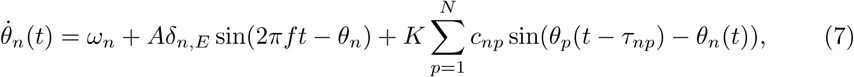

where *A* represents the amplitude of the external force and defines the coupling strength between the stimulus and the stimulated node. A fixed stimulation amplitude of *A* = 400 was used in all simulations, corresponding to the minimum value that ensured optimal synchronization of the stimulated node with the external signal, thereby maintaining consistent phase locking to the driving oscillation across all tested frequencies. Furthermore, *f* denotes the stimulation frequency in Hz and *E* identifies the node subjected to the external force, with *δ_n,E_* = 1 if *n* is equal to *E* and zero otherwise.

External stimulation was applied at the frequencies corresponding to the dominant peaks identified at the point of maximum spectral entropy during baseline (Fig 1). Stimulation with a sinusoidal oscillation of the form: *A* sin(2*πf* ), was applied separately to each network node, and independently at each of the frequency values *f* = 13.0, 29.4, 43.0 Hz. Stimulation frequencies were selected from the dominant peaks of the model’s spontaneous spectra (13.0, 29.4, and 43.0 Hz), corresponding to distinct oscillatory modes identified at maximal metastability. Notably, these frequencies align with well-established steady-state visual and auditory response bands—alpha (SSVEP, ∼10–13 Hz) and gamma (ASSR, ∼40 Hz) which enhances the biological plausibility of the model’s predictions.

Simulations were first performed using the parameters corresponding to the operating point of maximum entropy OP maxSE at baseline: {*K*, MD} = {4, 21 ms}, assuming that this coexistence of multiple oscillatory modes provides the ideal setting for illustrating the mechanism of frequency-dependent network entrainment.

Furthermore, given that this point of maximum spectral entropy is different from resting EEG power spectra as shown in Fig 1, we investigated how the model responds when the operating point is shifted to match the EEG power spectra OP EEG of different participants : {*K*, MD} = {5, 19 ms}, {5, 20 ms}, {5.5, 22 ms}, {4.5, 18 ms}, {4.5, 19 ms} and {5, 21 ms} (See S1 Fig), keeping the same stimulation frequencies.

### Measuring entrainment in the network model

A node is considered entrained to the stimulation frequency if its spectral power at that frequency is at least twice its baseline (no-stimulation) value, and if the phase-locking value (PLV) between the node and the driving signal, defined as the magnitude of the time-averaged complex phase difference between the two signals, 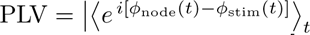 [54], exceeds 70%, indicating strong and consistent phase synchronization over time [55].

We further examined whether entrainment is influenced by the network’s structural properties. Specifically, we evaluated the linear relation between the number of entrained nodes and various network metrics, including weighted degree, betweenness centrality, degree centrality, closeness centrality, and connection delays (see S4 Fig).

Among these, weighted degree exhibited the strongest correlation with entrainment, suggesting that nodes with a higher total connection weight are more likely to facilitate the propagation of entrainment throughout the network.

### Simulations Design

We conducted a series of simulations to investigate how network entrainment is dependent on the stimulation Frequency, Location, and Operating Point.

More specifically, we explored how the system responds to external stimulation when changing:

- Stimulation Frequency and Target region

We first set the operating point of the model to maximize spectral entropy, where multiple oscillatory modes are detected in the power spectrum, and stimulated each brain region with a sinusoidal input at the peak frequency of each mode. In total 90 x 3 simulations were performed, with forced stimulation persisting for 300 seconds.

- Operating Points with distinct Power Spectra

We subsequently varied the operating point of the model by changing the global model parameters such that the simulated activity has a power spectrum that better matches the power spectra of EEG recordings from distinct participants.

Furthermore, we examined whether targeting sensory areas at specific frequencies produces patterns of neural activity consistent with those observed in empirical EEG recordings.

### Validation of the model

To assess frequency-selective correspondence between empirical and modeled entrainment, we compared EEG responses elicited at cortex-preferred drive frequencies with model predictions. Specifically, for visual entrainment, we used stimulation near each participant’s individual alpha frequency (IAF), and for auditory entrainment, we used 40 Hz stimulation; in both cases, we computed source-localized activation using the AAL90 parcellation at the driven frequency. These activation maps were then correlated with the model’s regional spectral profiles obtained under stimulation at each candidate frequency.

#### Dataset 1: Visual SSVEP (alpha entrainment)

Nineteen healthy adults (11 males, 8 females; 26 ± 3 years), with normal or corrected-to-normal vision and no history of neurological/psychiatric disorders, participated under ethics approval CEC170-18. Visual stimulation consisted of sinusoidally modulated light at each participant’s individual alpha frequency (IAF), delivered in blocks while the EEG was recorded. Analyses focused on frequency-resolved entrainment (spectral peaks at IAF and harmonics) and narrowband envelopes to quantify response magnitude and spread using the approach in [56].

#### Dataset 2: Auditory ASSR (gamma entrainment)

An independent cohort of 13 healthy adults (6 males, 7 females; 25 ± 2 years) completed an auditory steady-state response protocol, approved under ethics code CEC230-21. Stimulation consisted of 40-Hz amplitude-modulated tones (1-kHz carrier). The protocol comprised five blocks, each with 20 trials of 5-s stimulation. We quantified ASSR amplitude from narrowband (around 40 Hz) envelopes, providing a benchmark for model-based predictions in the auditory cortex.

To compare the model results with the empirical visual and auditory stimulation data, we analyzed the brain regions that showed significant activation during real stimulation and those exhibiting entrainment in the model simulations. In the empirical data, stimulation was bilateral, whereas in the simulations it was applied focally to a single primary cortical node: the left Calcarine cortex for the visual condition and the left Heschl’s gyrus for the auditory condition. Despite this difference, this approach allows assessing the model’s ability to reproduce the frequency selectivity observed experimentally, that is, whether the strongest responses occur at frequencies closest to those used in the empirical data.

To evaluate the correspondence between simulated and empirical spatial entrainment patterns, we used two metrics: Pearson’s correlation, which captures the linear association between entrainment intensities, and cosine similarity, which quantifies the spatial correspondence of the entrainment patterns regardless of their magnitude. In both cases, the analysis was based on vectors representing the activation magnitude of each brain region. For the empirical data, activation magnitude was obtained by averaging voxel-wise activation values within each brain region, whereas for the simulated data, the regional activation was used directly.

## Results

### Network Entrainment depends on Stimulation Frequency and Target region

We stimulated each of the model’s 90 nodes individually (a node corresponds to a brain region in the AAL90 parcellation [48]) at each of the peak frequencies of the three more prominent oscillatory modes (13, 29.4, and 43 Hz) identified in the operating point of maximum metastability OP maxSE as quantified by spectral entropy [11](see yellow square in Fig 1). We use the model coupling and delay parameters pair {*K*, MD}, also corresponding to the OP maxSE.

We quantified stimulation spread as the number of entrained nodes and correlated it with graph-theoretical metrics of the stimulated region (weighted degree, degree, closeness, betweenness, and delay). Weighted degree showed the strongest association with the number of entrained nodes indicating that this structural property is a robust predictor of regional susceptibility to stimulation. Full correlation results are provided in S4 Fig.

For subsequent analyses, we selected three representative nodes spanning the range of weighted degree: right precuneus (highest), left precentral gyrus (intermediate), and right paracentral lobule (lowest). Across all three sites, periodic stimulation produced entrainment that extended beyond the stimulated region, with the spatial spread depending on both stimulation frequency and stimulation site as is shown in Fig 2.

**Fig 2.**
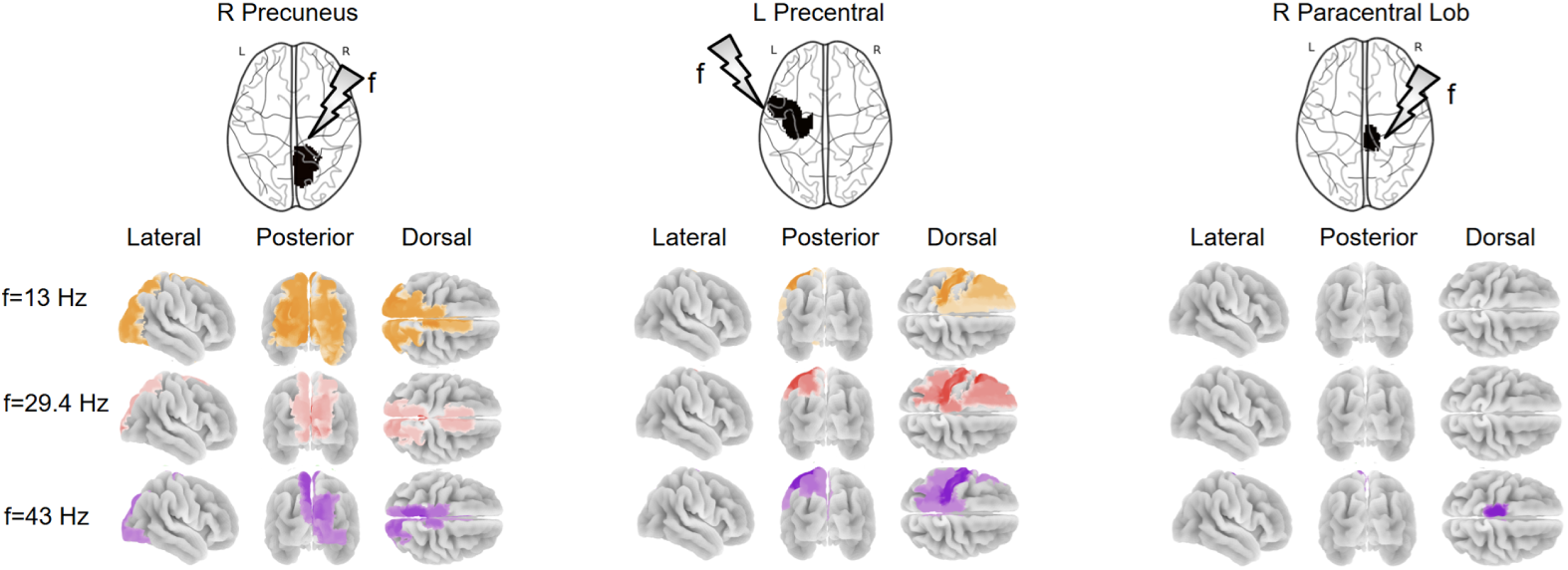
Selective network entrainment by driving frequency and stimulation site. Functional networks that synchronise to the driving force depend not just on the node to which the stimulation is applied, but also on the driving frequency, such that selective network modulation could be achieved by the choice of stimulation parameters. The weighted degree of a node (total sum of connection weights) affects the extension of entrainment (stimulation spread). Cortical nodes with high (right precuneus), intermediate (left precentral gyrus), or low (right paracentral lobule) weighted degree were stimulated at each of the five oscillatory frequencies emerging at the maximum spectral entropy. Nodes exhibiting entrainment are shown for each stimulation site and frequency, highlighting distinct spatial propagation patterns. These findings demonstrate that functional networks synchronized to the driving force depend on both the stimulation frequency and the targeted site, indicating that selective network modulation can be achieved through proper tuning of stimulation parameters.

As expected, stimulating the right precuneus—the node with the highest weighted degree—recruited the widest network, exceeding the spread observed for the precentral gyrus and paracentral lobule. At lower stimulation frequencies, stimulation of the right precuneus led to bilateral propagation of entrainment into occipital, dorsomedial parietal, and medial frontal cortices; the spatial extent of entrainment monotonically decreased as stimulation frequency increased.

Stimulating the precentral gyrus also engaged frequency-specific subnetworks, but the dependence on frequency was weaker, yielding less systematic propagation patterns. In contrast, the paracentral lobule showed no entrainment at low frequencies; oscillatory responses emerged only above 40 Hz and remained local without propagation. Detailed of the regions and the frequency-by-frequency outcomes at the operating point of maximum entropy are shown in S5 Table, S6 Fig, S7 Fig and S8 Fig.

We performed two complementary analyses targeting distinct aspects of neural entrainment. In the first, we analyzed whether synchrony at the driving frequency could propagate through the network, entraining nodes beyond those directly connected to the stimulated region (”entrainment propagation”, S9 Fig shows the AAL90 structural connection map). In the second, we investigated whether stimulation at specific frequencies could effectively engage deep cortical and subcortical structures, even when applied to superficial cortical nodes. Specifically, we examined entrainment propagation at 13, 29.4, and 43 Hz using the parameter set corresponding to OP maxSE.

S10 Fig shows that frequency-dependent entrainment can reach remote nodes not structurally connected to the stimulation site, suggesting that rhythmic input spreads through indirect pathways mediated by intermediary nodes. S11 Fig shows that frequency-dependent stimulation successfully recruits deep brain areas, indicating that rhythmic input can influence subcortical targets through cortical stimulation. Overall results show that neural entrainment depends on both the stimulation frequency and network topology. This frequency selectivity aligns with prior research showing that lower-frequency oscillations facilitate long-range communication, while higher frequencies tend to remain more localized within cortical circuits [46, 57, 58].

### Network Entrainment depends on Operating Point

The variability observed in the spectral features of real EEG recordings is captured in our model by changes in {*K*, MD} pairs. To investigate how these differences in operating point influence network responsiveness, we examined the spread of entrainment as a function of stimulation frequency for different operating points, using the right precuneus (the node with the maximum weighted degree) as the stimulation site. The {*K*, MD} pairs were chosen to reflect those associated with EEG spectra from resting healthy individuals, as illustrated in S1 Fig.

Using {*K*, MD} pairs distinct from the one associated with maximum metastability reproduced the main finding, namely, that the broadest spread of entrainment consistently occurred at the lowest stimulation frequency, regardless of the selected {*K*, MD} combination (Fig 3). Furthermore, each stimulation frequency entrained a core subnetwork, which remained stable across {*K*, MD} combinations. At a stimulation frequency of 13 Hz, this core subnetwork comprised the majority of the occipital brain regions along with large portions of the parietal, cingulate, and limbic cortices in both hemispheres. When the stimulation frequency increased to 29.4 Hz, the core subnetwork was reduced to the bilateral precuneus, the right superior parietal lobe, and the right middle cingulum. At 43 Hz, only the right precuneus remained consistently activated across all {*K*, MD} pairs. The spread of entrainment beyond these core subnetworks was modulated by the {*K*, MD} parameters, as different combinations led to the recruitment of additional nodes and the disengagement of previously participating brain regions as illustrated in Fig 3. Nodes recruited by stimulation included deep cortical brain regions and subcortical structures.

**Fig 3.**
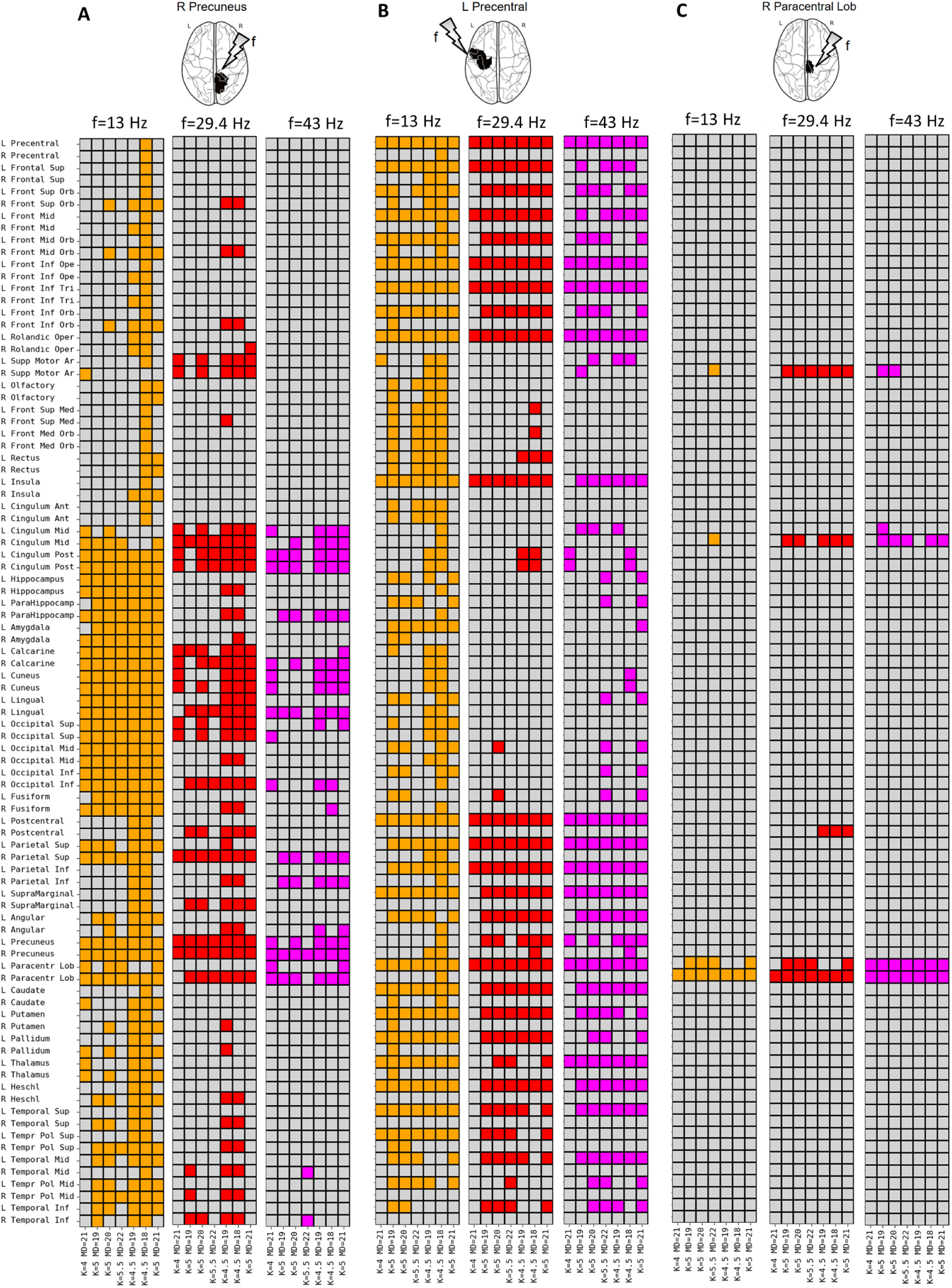
Network entrainment depends on the driving frequency and operating point of the system. Effects of 13 Hz, 29.4 Hz, and 43 Hz stimulation of the right precuneus (A), left precentral gyrus (B) and right paracentral lobule (C), when simulating with different working points ( see S1 Fig. The entrainment is illustrated by coloring the nodes that synchronize with the driving frequency. While there is a consistent core network entrained by each distinct driving frequency, the engagement of some nodes depends on the operating point.

### Frequency-specific entrainment of sensory systems

To validate the network entrainment results, we examined whether rhythmic stimulation of sensory cortical nodes in the model could reproduce the frequency-dependent patterns observed in electroencephalographic steady-state evoked potentials (SSEPs) during visual and auditory stimulation [56, 59–61]. Specifically, stimulation was applied to the left Calcarine cortex and Heschl’s gyrus, representing the primary cortical targets of visual and auditory inputs at frequencies near their canonical resonances (13 Hz for the visual network and 43 Hz for the auditory network). We evaluated the stimulation across two operating points of the model, encompassing the maximum-metastability regime and the operating point derived of the mean of the participants.

Fig 4 shows modality-dependent frequency preferences consistent with prior EEG studies of sensory processing at two operating points: the OP maxSE and the OP EEG. In the case of the visual cortex (left Calcarine), entrainment was reliably induced across all tested stimulation frequencies and was consistently observed across both {*K*, MD} combinations. As anticipated, entrainment propagated beyond the stimulation site, often extending to the ipsilateral lateral parietal and temporal cortices, as well as to the contralateral occipital lobe, depending on stimulation frequency and the operating point employed in the simulations. It is noteworthy that for every operating point , the strongest entrainment was elicited by 13-Hz stimulation.

**Fig 4.**
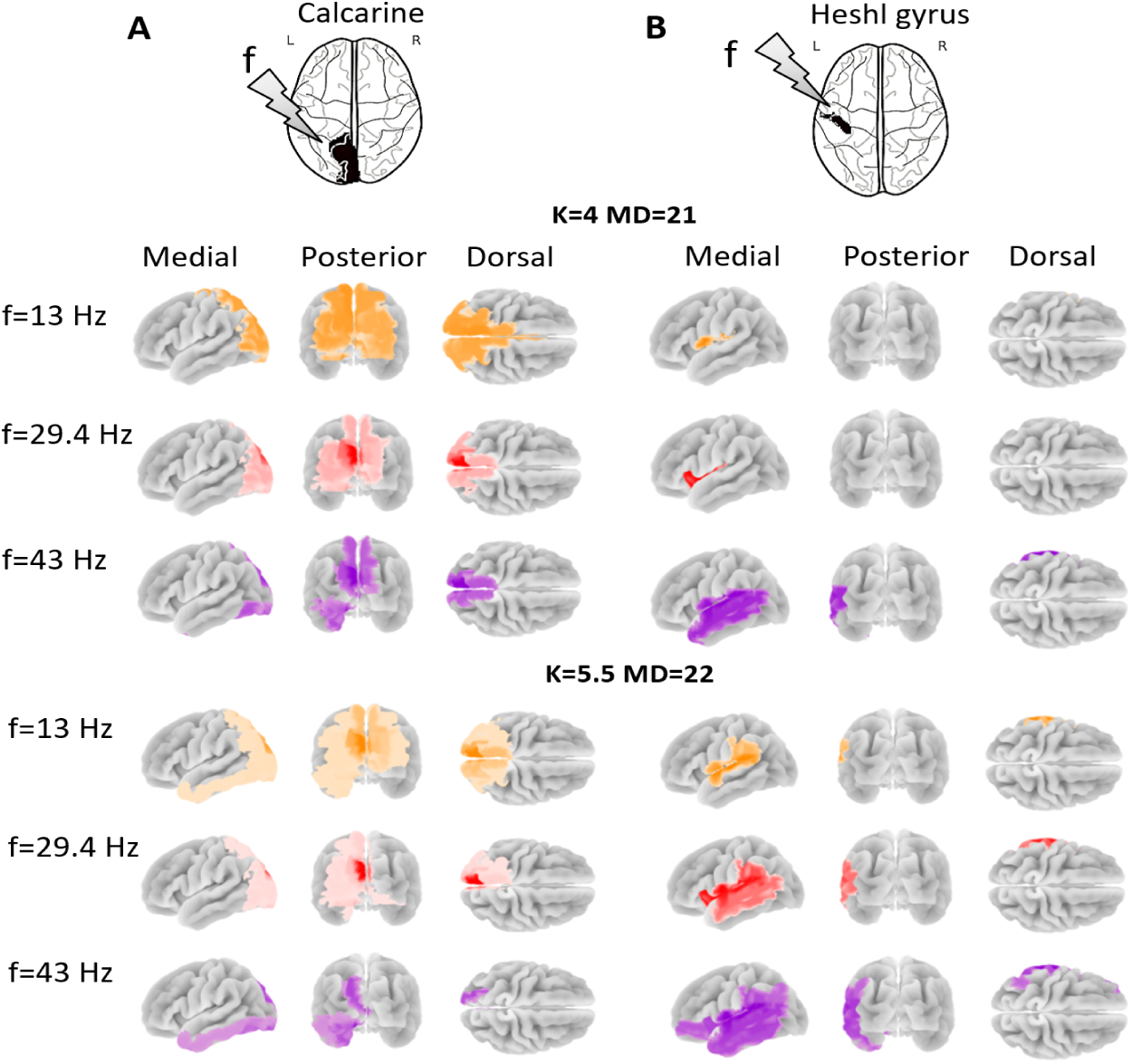
Entrainment effects across sensory cortices as a function of stimulation frequency and operating point{*K*, MD}. A. Left Calcarine (Visual Cortex): Strongest entrainment at 13 Hz, aligning with alpha-band oscillations known to regulate visual processing and attentional gating. B. Left Heschl’s Gyrus (Auditory Cortex): Preferred entrainment at 43 Hz, consistent with gamma-band dominant frequency in auditory networks, reflecting the temporal precision required for auditory processing. These findings underscore the role of intrinsic oscillatory dynamics in shaping modality-specific entrainment and demonstrate how neuromodulation effects depend on both regional specialization and individual variability in network state.

Similarly, stimulation of the auditory cortex (left Heschl’s Gyrus) resulted in robust entrainment at all tested frequencies, exhibiting clear frequency-specific patterns. Across both {*K*, MD} combinations, the most pronounced entrainment was observed at a stimulation frequency of 43 Hz. The broadest spatial propagation of entrainment occurred under simulation parameters of the OP EEG,encompassing the entire temporal lobe, as well as the lateral parietal and inferior frontal regions. The entrainment also propagated to ipsilateral regions of the stimulated hemisphere.

To assess the model results, we compared the effects of stimulation in the primary visual and auditory cortex nodes with real bilateral visual and auditory stimulation data. For visual stimulation, subjects were stimulated at the alpha frequency, while for auditory stimulation, they received stimulation at 40 Hz (see Methods section ). Although stimulation in the model was restricted to a single node, the primary objective of this validation was to assess the model’s capacity to reproduce frequency-selective entrainment. Accordingly, simulations were conducted at three stimulation frequencies (13, 29.4, and 43 Hz).

S12 Fig and S13 Fig illustrate the spatial patterns of entrainment obtained from the simulated and empirical datasets, while S14 Table and S15 Table present the quantitative measures of correspondence across all stimulation frequencies. The best agreement between the model and empirical data was obtained for visual stimulation at 13 Hz, where 16 of the 36 activated regions in the empirical data coincided with those predicted by the model, yielding a Pearson correlation of 0.40 (*p_value_ <* 0.05) and a cosine similarity of 0.50. For auditory stimulation, the best results were found at 43 Hz, where 9 of the 39 activated regions in the empirical data overlapped with those predicted by the model, yielding a Pearson correlation of 0.37 (*p_value_ <* 0.05) and a cosine similarity of 0.43.

## Discussion

### Modeling brain networks as a bridge between mechanistic understanding of oscillatory dynamics and and clinical neurorehabilitation

Neurorehabilitation through non-invasive neurostimulation has gained increasing attention as a means of restoring or enhancing brain function following neurological injury or disease. Evidence from clinical studies has shown that rhythmic stimulation can modulate oscillatory brain activity and improve behavioural outcomes in several patient populations. Although numerous neurostimulation studies have demonstrated promising results, standardized protocols have yet to be established, and outcomes still differ considerably across individuals, stimulation sites, and frequencies. This variability suggests that uniform stimulation protocols are unlikely to achieve optimal effects, particularly during clinically critical phases, such as the acute period following stroke, when direct testing in patients is often not possible. In this context, modeling brain network dynamics based on real structural connectivity information provides a controlled framework for systematically probing the mechanisms underlying neurostimulation, helping to bridge the gap between theoretical understanding of the brain oscillatory dynamics and clinical application. By working in silico, it becomes possible to explore a large parameter space systematically and ethically, during phases of heightened plasticity when direct patient intervention is constrained, enabling systematic exploration of stimulation parameters beyond what can be tested experimentally.

### Whole-brain network models reveal site-dependent propagation of entrainment

We used a whole-brain network model of coupled Kuramoto oscillators, constrained by empirical human structural connectivity, to examine how stimulation frequency, target site, and network operating point interact to shape large-scale brain dynamics.

The simulations demonstrated that the structural embedding of the stimulation site was a determinant of spatial extent of entrainment. Stimulation of hub regions with high weighted degree, such as the precuneus, produced broad bilateral propagation across parietal, occipital, and medial frontal regions, while stimulation of less connected regions resulted in more localized effects. Quantitative analyses confirmed that weighted degree best predicted the number of entrained nodes, consistent with the idea that strongly connected hubs act as effective broadcast points, supporting widespread network engagement. This finding aligns with empirical neuroimaging studies showing that stimulating the precuneus can elicit widespread cortical responses and modulate large-scale network dynamics (Koch et al., 2022). By directly linking stimulation spread to structural network metrics, the model provided mechanistic grounding for site-specific variability observed in empirical neuromodulation outcomes.

### Frequency-dependent entrainment reveals structural filtering of rhythmic inputs

Beyond stimulation site, our results demonstrate that stimulation frequency is another major determinant of network response. The network model showed distinct frequency-selective entrainment patterns, with lower frequencies supporting broad, long-range propagation and higher frequencies producing more localized effects.

This frequency dependence reflected the interaction between external rhythmic inputs and the network’s structural and dynamical properties: low-frequency rhythms are more robust to transmission delays and thus better suited for global communication, whereas higher frequency oscillations tend to remain localized due to phase dispersion across longer connections [5, 58, 62, 63]. These results are consistent with theoretical work on resonance and signal transmission in delay-coupled oscillator networks, which predicts frequency-dependent filtering by the network structure [8]. Importantly, different frequencies engaged different subnetworks even when stimulating the same region, indicating that frequency can be used as a control parameter to select specific communication channels within the network. This provides a mechanistic explanation for the empirically observed frequency tuning of stimulation effects (e.g., SSVEP [32] and ASSR [61]), and supports the view that effective stimulation depends not only on where but also on how fast we stimulate.

### Global network operating regimes modulate the impact of stimulation

In addition to stimulation parameters, the global operating point of the model, determined by coupling strength and mean conduction delay {*K*, MD}, modulated both the extent and selectivity of entrainment. Shifting the model parameters to match individual EEG-derived spectra altered how susceptible the network was to the external driving and how far entrainment propagated. These results suggest that inter-individual differences in network dynamics, reflected in global parameters, can significantly influence stimulation outcomes even with identical structural connectivity and stimulation parameters. This may partly explain the variability observed in clinical and experimental studies, where some individuals exhibit strong, widespread entrainment, while others show weak or absent responses. Modeling provides a way to estimate and incorporate such individual operating regimes before stimulation is applied, guiding frequency and site selection on a patient-specific basis. Such connectome-informed neuromodulation may enhance the precision and reproducibility of interventions such as tACS or rTMS, bringing personalized neurostimulation strategies closer to clinical reality.

### Empirical validation supports biologically plausible frequency selectivity

To assess the biological plausibility of the model, we compared its predictions with empirical EEG responses obtained during rhythmic sensory stimulation. Stimulation of the visual cortex at 13 Hz and of the auditory cortex at 43 Hz reproduced the characteristic spatial activation patterns of steady-state visual and auditory responses (SSVEP and ASSR, respectively) observed in real data [56, 59–61]. This correspondence indicates that the model captured not only generic principles of network synchronization but also physiologically meaningful, frequency-specific entrainment. These findings support the view that connectome-based modeling provides a suitable framework for exploring and predicting large-scale neural responses under controlled stimulation conditions.

Quantitative validation revealed modest yet significant spatial correlations between simulated and empirical activation maps (Pearson *r* = 0.40 for visual and *r* = 0.37 for auditory stimulation; both *p <* 0.05). Importantly, these peak correlations were obtained precisely at the stimulation frequencies that matched the preferred response bands observed in the empirical data—13 Hz for visual entrainment in the alpha range and 43 Hz for auditory entrainment in the gamma range. This alignment reinforces the model’s ability to reproduce frequency-selective resonance consistent with known sensory dynamics. The observed discrepancies between simulated and empirical maps likely stem from simplifications inherent in the current framework, including unilateral stimulation, reliance on a cortex-only connectome, and the omission of long-range subcortical and interhemispheric pathways that contribute to multisensory communication. Additional variability in the EEG-derived maps, due to individual anatomical differences, tissue conductivity, and source reconstruction uncertainty, may also contribute to the reduced spatial correspondence.

### Limitations ans future directions

Several limitations of this study should be acknowledged. First, the model relied exclusively on cortical structural connectivity, omitting subcortical and long-range interhemispheric tracts that are known to mediate information flow between auditory and visual pathways. The absence of these deep and cross-hemispheric connections likely constrained the model’s ability to reproduce the full spatial extent of sensory entrainment observed in EEG data. Second, stimulation in the model was applied unilaterally to a single cortical hemisphere, whereas the empirical protocols involved bilateral sensory stimulation. This asymmetry may have underestimated inter-hemispheric propagation effects, contributing to the reduced spatial overlap between simulated and empirical activation patterns. Third, the model represented large-scale dynamics using phase-coupled oscillators with homogeneous intrinsic frequencies—a simplification that neglects regional heterogeneity, non-linear interactions, and cross-frequency coupling, all of which can substantially shape steady-state responses and network-level synchronization.

In addition, the use of a group-level connectome does not account for subject-specific anatomical variability that may influence stimulation propagation and frequency sensitivity, potentially contributing to the modest correlations observed during model validation.

Future work should aim to address these limitations by incorporating individualized structural and functional connectomes, as well as bilateral stimulation protocols, which could enhance the model’s ability to replicate empirical EEG topographies and improve its predictive accuracy. Expanding the framework to simulate plasticity mechanisms and transient network dynamics would also allow exploration of long-term adaptations to stimulation, bridging toward clinically relevant timescales. Finally, integrating closed-loop adaptive stimulation strategies within these computational models could enable real-time optimization of stimulation parameters based on predicted network responses.

By addressing these aspects, the proposed framework could evolve into a robust predictive tool for connectome-informed neuromodulation, capable of guiding personalized stimulation design and parameter tuning—particularly valuable in acute and early post-injury phases where empirical experimentation remains limited.

## Conclusion

This study demonstrates that frequency-selective network entrainment naturally emerges from the interplay between the brain’s structural connectivity and its intrinsic oscillatory dynamics. Using a connectome-based model of coupled Kuramoto oscillators, we systematically showed how stimulation frequency, target site, and global operating state jointly determine large-scale synchronization patterns. The model revealed that stimulation propagates across distributed subnetworks rather than remaining confined to the target region, following the topological organization of the connectome. Low-frequency inputs favored widespread, long-range synchronization, while higher frequencies produced more localized responses—an inverse relationship consistent with empirical observations from steady-state visual and auditory responses.

By reproducing modality-specific frequency preferences observed in empirical EEG data—alpha-band entrainment for visual and gamma-band entrainment for auditory stimulation—the model captured physiologically meaningful, frequency-dependent activation patterns. These results highlight how network structure constrains stimulation effects and provide mechanistic insight into frequency-selective entrainment across sensory systems. Beyond explaining observed phenomena, this framework establishes a foundation for predictive, connectome-informed neuromodulation. Future extensions integrating individualized connectomes, more biophysically detailed models, and adaptive closed-loop control could enable the rational design of personalized stimulation protocols, ultimately improving the precision and efficacy of non-invasive brain stimulation in both research and clinical contexts.

## Supporting information

**S1 Fig.**
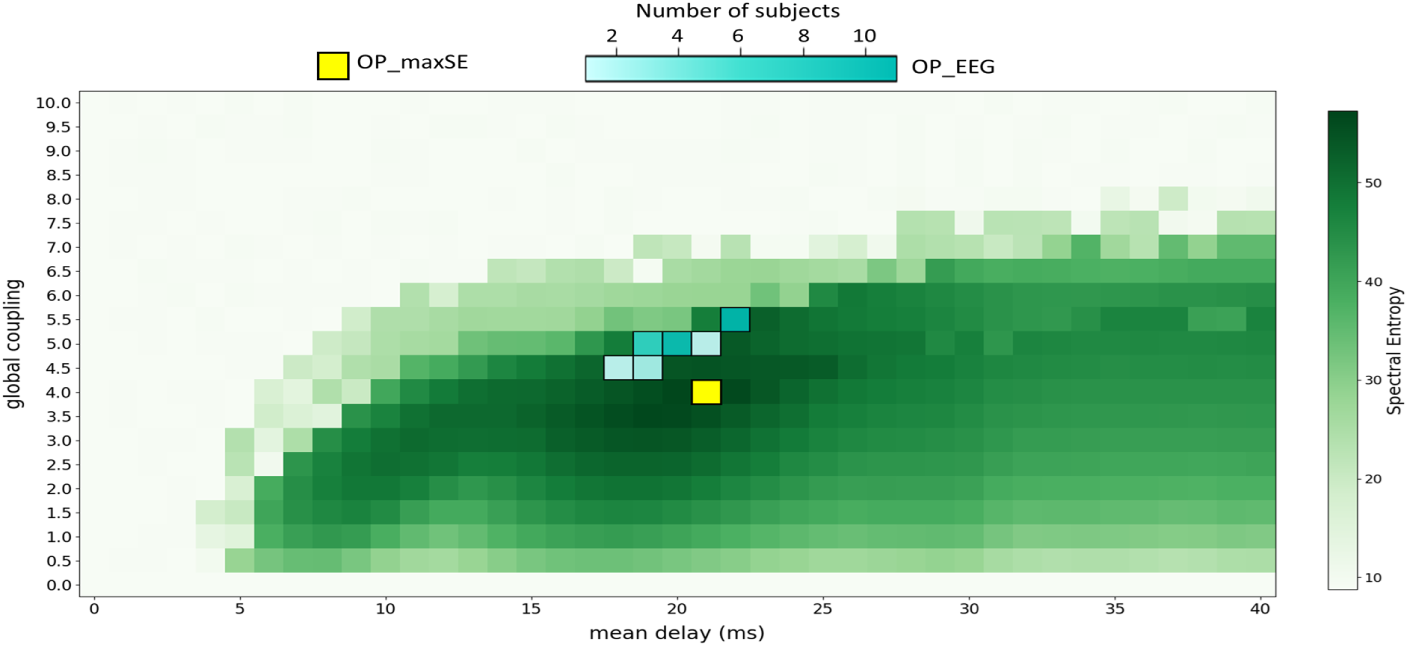
Spectral entropy in the parameter space exploration of global coupling (K) and mean delay (MD). The yellow square highlights the maximun value of spectral entropy. The blue squares represents de location of the fitting real EEG data.

**S2 Fig.**
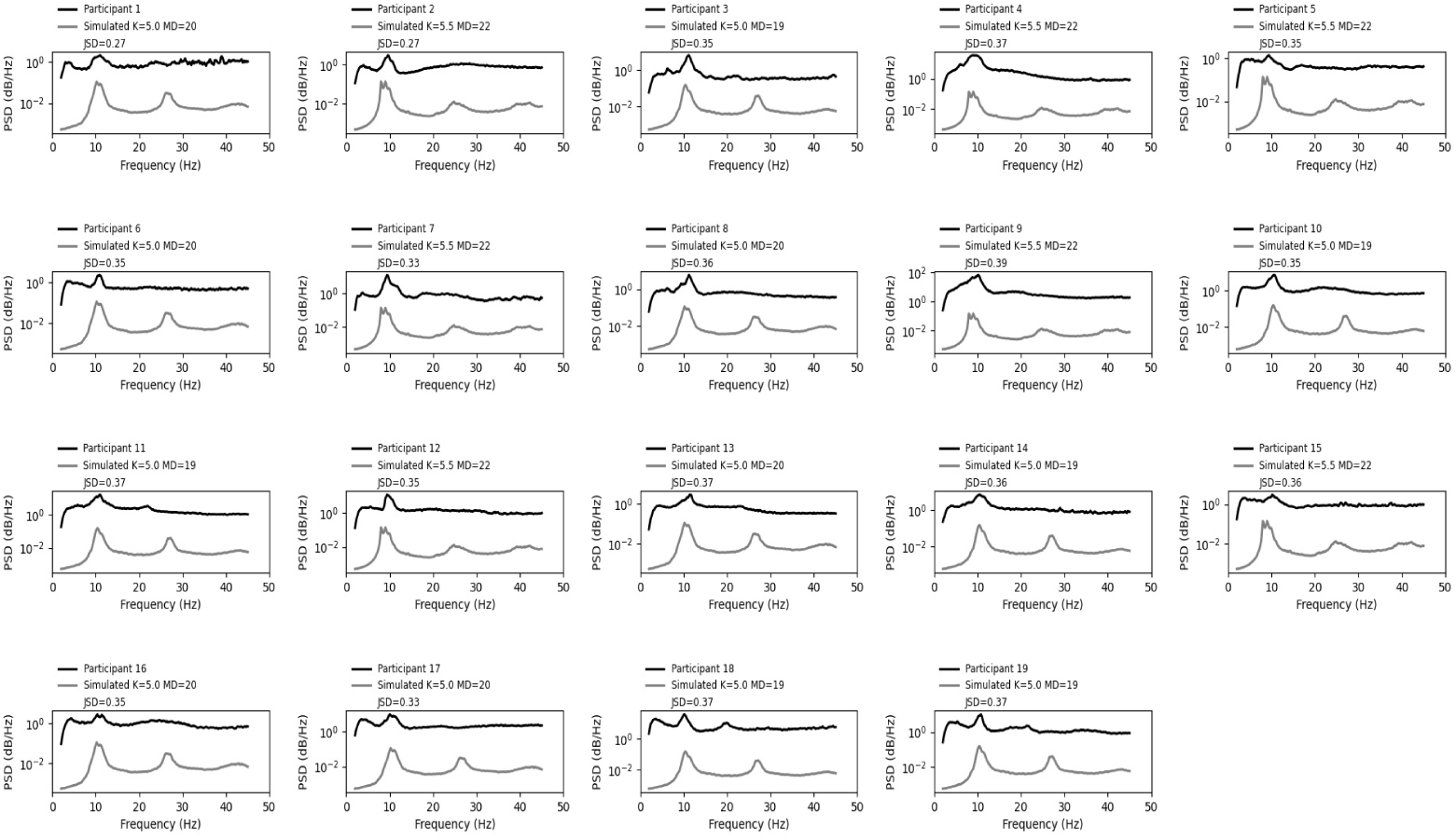
Mapping the spectrum from the kuramoto model to the spectrum of real EEG subjects participated under ethics approval CEC170-18. It shows the spectrum of the Kuramoto model (marked in gray) that had the smallest Jensen-Shannon (JS) distance compared to the real EEG spectra of the subjects (marked in black).

**S3 Fig.**
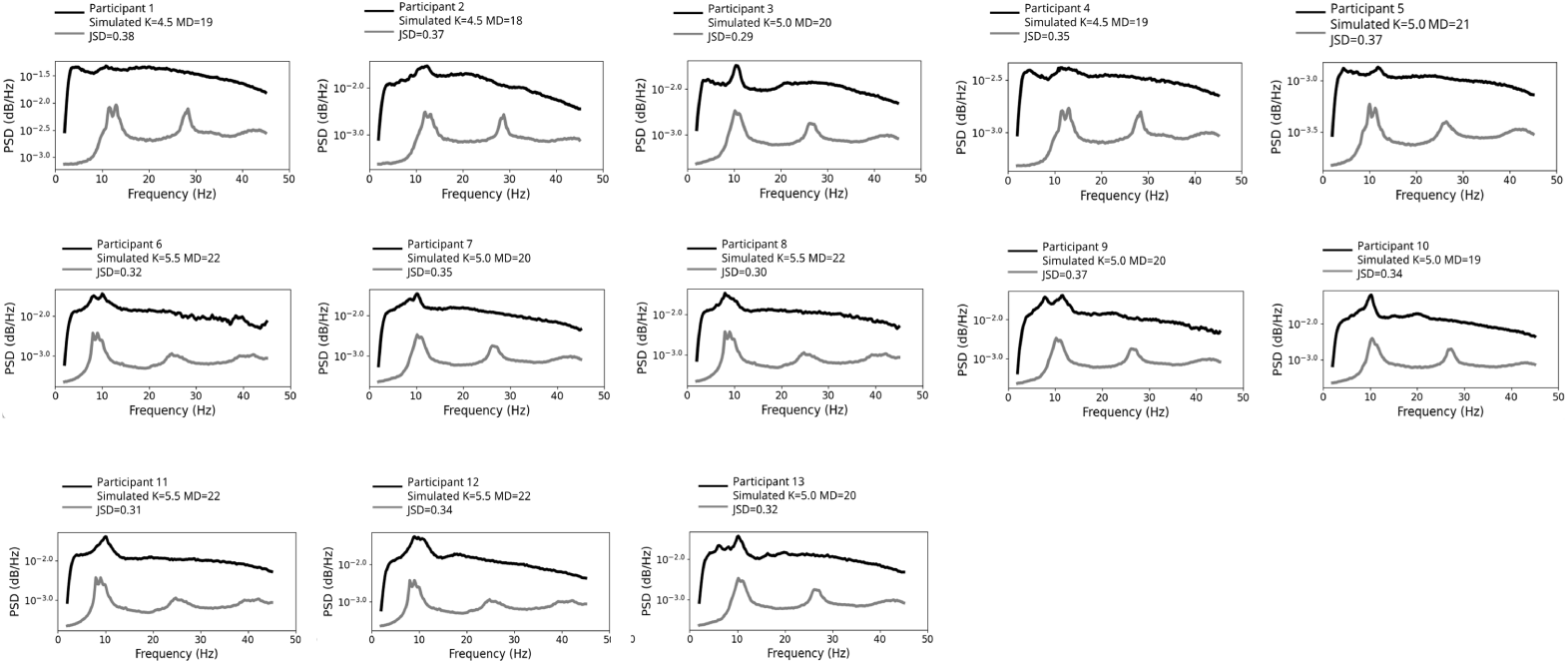
Mapping the spectrum from the kuramoto model to the spectrum of real EEG subjects participated under ethics approval CEC230-21. It shows the spectrum of the Kuramoto model (marked in gray) that had the smallest Jensen-Shannon (JS) distance compared to the real EEG spectra of the subjects (marked in black).

**S4 Fig.**
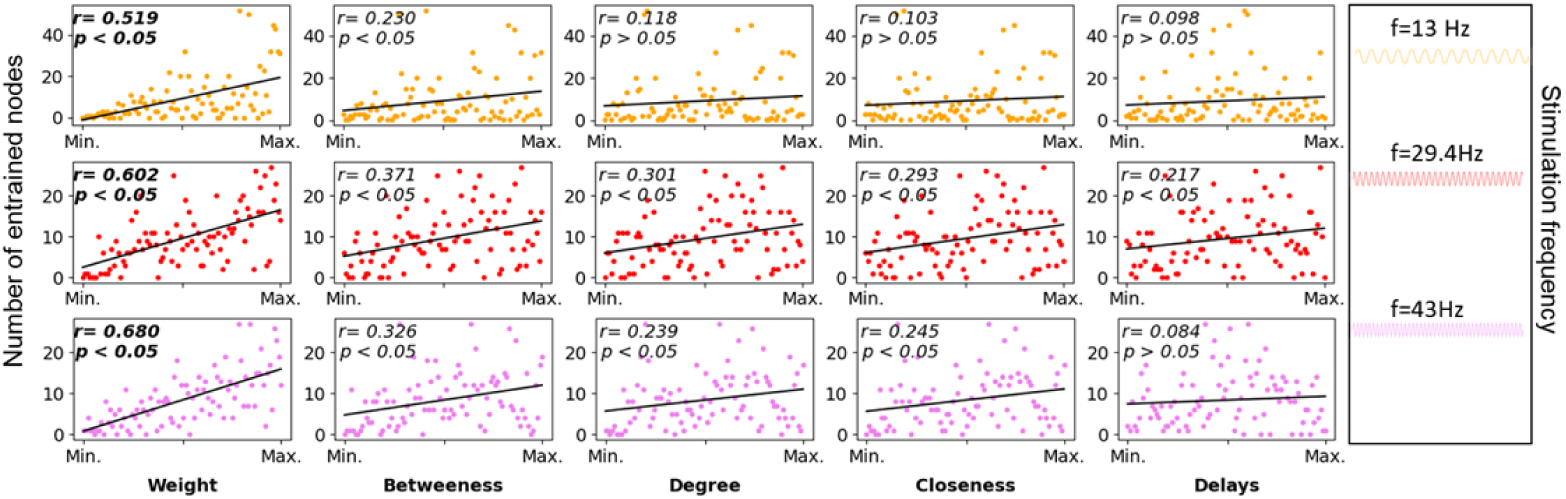
Stimulation effects across nodes at different frequencies, sorted by structural features. These features describe key aspects of the network topology, including connection strength (weighted degree), number of direct links (degree), efficiency in reaching other nodes (closeness), control over information flow paths (betweenness), and communication time between regions (delays). Each column displays nodes in descending order based on the corresponding feature, along with a linear fit. Across all frequencies, node weighted degree showed the strongest linear association with stimulation effects.

**S5 Table.**
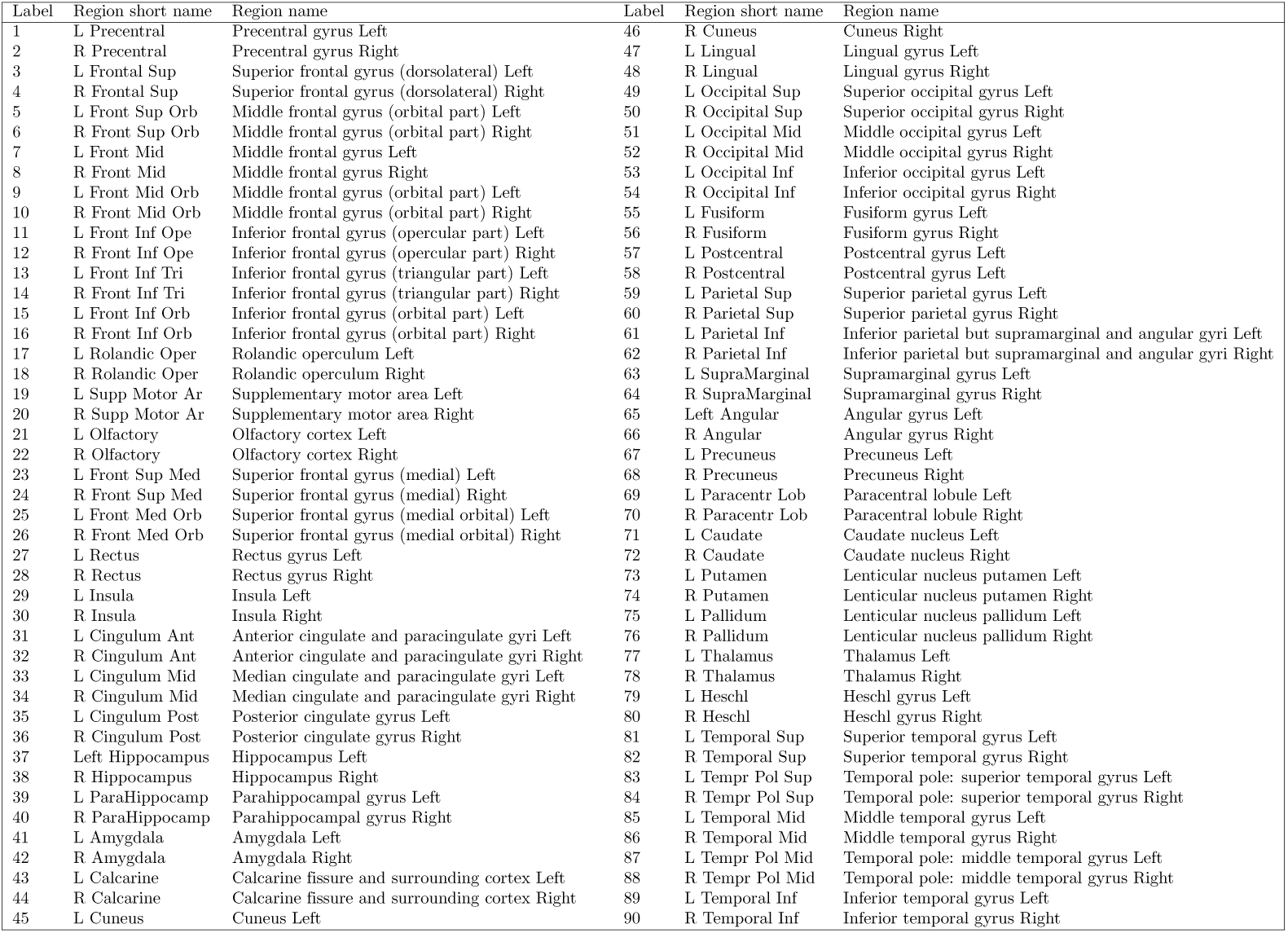
Glossary of AAL90 regions.

| Label | Region short name | Region name | Label | Region short name | Region name |
| --- | --- | --- | --- | --- | --- |
| 1 | L Precentral | Precentral gyrus Left | 46 | R Cuneus | Cuneus Right |
| 2 | R Precentral | Precentral gyrus Right | 47 | L Lingual | Lingual gyrus Left |
| 3 | L Frontal Sup | Superior frontal gyrus (dorsolateral) Left | 48 | R Lingual | Lingual gyrus Right |
| 4 | R Frontal Sup | Superior frontal gyrus (dorsolateral) Right | 49 | L Occipital Sup | Superior occipital gyrus Left |
| 5 | L Front Sup Orb | Middle frontal gyrus (orbital part) Left | 50 | R Occipital Sup | Superior occipital gyrus Right |
| 6 | R Front Sup Orb | Middle frontal gyrus (orbital part) Right | 51 | L Occipital Mid | Middle occipital gyrus Left |
| 7 | L Front Mid | Middle frontal gyrus Left | 52 | R Occipital Mid | Middle occipital gyrus Right |
| 8 | R Front Mid | Middle frontal gyrus Right | 53 | L Occipital Inf | Inferior occipital gyrus Left |
| 9 | L Front Mid Orb | Middle frontal gyrus (orbital part) Left | 54 | R Occipital Inf | Inferior occipital gyrus Right |
| 10 | R Front Mid Orb | Middle frontal gyrus (orbital part) Right | 55 | L Fusiform | Fusiform gyrus Left |
| 11 | L Front Inf Ope | Inferior frontal gyrus (opercular part) Left | 56 | R Fusiform | Fusiform gyrus Right |
| 12 | R Front Inf Ope | Inferior frontal gyrus (opercular part) Right | 57 | L Postcentral | Postcentral gyrus Left |
| 13 | L Front Inf Tri | Inferior frontal gyrus (triangular part) Left | 58 | R Postcentral | Postcentral gyrus Right |
| 14 | R Front Inf Tri | Inferior frontal gyrus (triangular part) Right | 59 | L Parietal Sup | Superior parietal gyrus Left |
| 15 | L Front Inf Orb | Inferior frontal gyrus (orbital part) Left | 60 | R Parietal Sup | Superior parietal gyrus Right |
| 16 | R Front Inf Orb | Inferior frontal gyrus (orbital part) Right | 61 | L Parietal Inf | Inferior parietal but supramarginal and angular gyri Left |
| 17 | L Rolandic Oper | Rolandic operculum Left | 62 | R Parietal Inf | Inferior parietal but supramarginal and angular gyri Right |
| 18 | R Rolandic Oper | Rolandic operculum Right | 63 | L SupraMarginal | Supramarginal gyrus Left |
| 19 | L Supp Motor Ar | Supplementary motor area Left | 64 | R SupraMarginal | Supramarginal gyrus Right |
| 20 | R Supp Motor Ar | Supplementary motor area Right | 65 | Left Angular | Angular gyrus Left |
| 21 | L Olfactory | Olfactory cortex Left | 66 | R Angular | Angular gyrus Right |
| 22 | R Olfactory | Olfactory cortex Right | 67 | L Precuneus | Precuneus Left |
| 23 | L Front Sup Med | Superior frontal gyrus (medial) Left | 68 | R Precuneus | Precuneus Right |
| 24 | R Front Sup Med | Superior frontal gyrus (medial) Right | 69 | L Paracentr Lob | Paracentral lobule Left |
| 25 | L Front Med Orb | Superior frontal gyrus (medial orbital) Left | 70 | R Paracentr Lob | Paracentral lobule Right |
| 26 | R Front Med Orb | Superior frontal gyrus (medial orbital) Right | 71 | L Caudate | Caudate nucleus Left |
| 27 | L Rectus | Rectus gyrus Left | 72 | R Caudate | Caudate nucleus Right |
| 28 | R Rectus | Rectus gyrus Right | 73 | L Putamen | Lenticular nucleus putamen Left |
| 29 | L Insula | Insula Left | 74 | R Putamen | Lenticular nucleus putamen Right |
| 30 | R Insula | Insula Right | 75 | L Pallidum | Lenticular nucleus pallidum Left |
| 31 | L Cingulum Ant | Anterior cingulate and paracingulate gyri Left | 76 | R Pallidum | Lenticular nucleus pallidum Right |
| 32 | R Cingulum Ant | Anterior cingulate and paracingulate gyri Right | 77 | L Thalamus | Thalamus Left |
| 33 | L Cingulum Mid | Median cingulate and paracingulate gyri Left | 78 | R Thalamus | Thalamus Right |
| 34 | R Cingulum Mid | Median cingulate and paracingulate gyri Right | 79 | L Heschl | Heschl gyrus Left |
| 35 | L Cingulum Post | Posterior cingulate gyrus Left | 80 | R Heschl | Heschl gyrus Right |
| 36 | R Cingulum Post | Posterior cingulate gyrus Right | 81 | L Temporal Sup | Superior temporal gyrus Left |
| 37 | Left Hippocampus | Hippocampus Left | 82 | R Temporal Sup | Superior temporal gyrus Right |
| 38 | R Hippocampus | Hippocampus Right | 83 | L Tempr Pol Sup | Temporal pole: superior temporal gyrus Left |
| 39 | L ParaHippocamp | Parahippocampal gyrus Left | 84 | R Tempr Pol Sup | Temporal pole: superior temporal gyrus Right |
| 40 | R ParaHippocamp | Parahippocampal gyrus Right | 85 | L Temporal Mid | Middle temporal gyrus Left |
| 41 | L Amygdala | Amygdala Left | 86 | R Temporal Mid | Middle temporal gyrus Right |
| 42 | R Amygdala | Amygdala Right | 87 | L Tempr Pol Mid | Temporal pole: middle temporal gyrus Left |
| 43 | L Calcarine | Calcarine fissure and surrounding cortex Left | 88 | R Tempr Pol Mid | Temporal pole: middle temporal gyrus Right |
| 44 | R Calcarine | Calcarine fissure and surrounding cortex Right | 89 | L Temporal Inf | Inferior temporal gyrus Left |
| 45 | L Cuneus | Cuneus Left | 90 | R Temporal Inf | Inferior temporal gyrus Right |

**S6 Fig.**
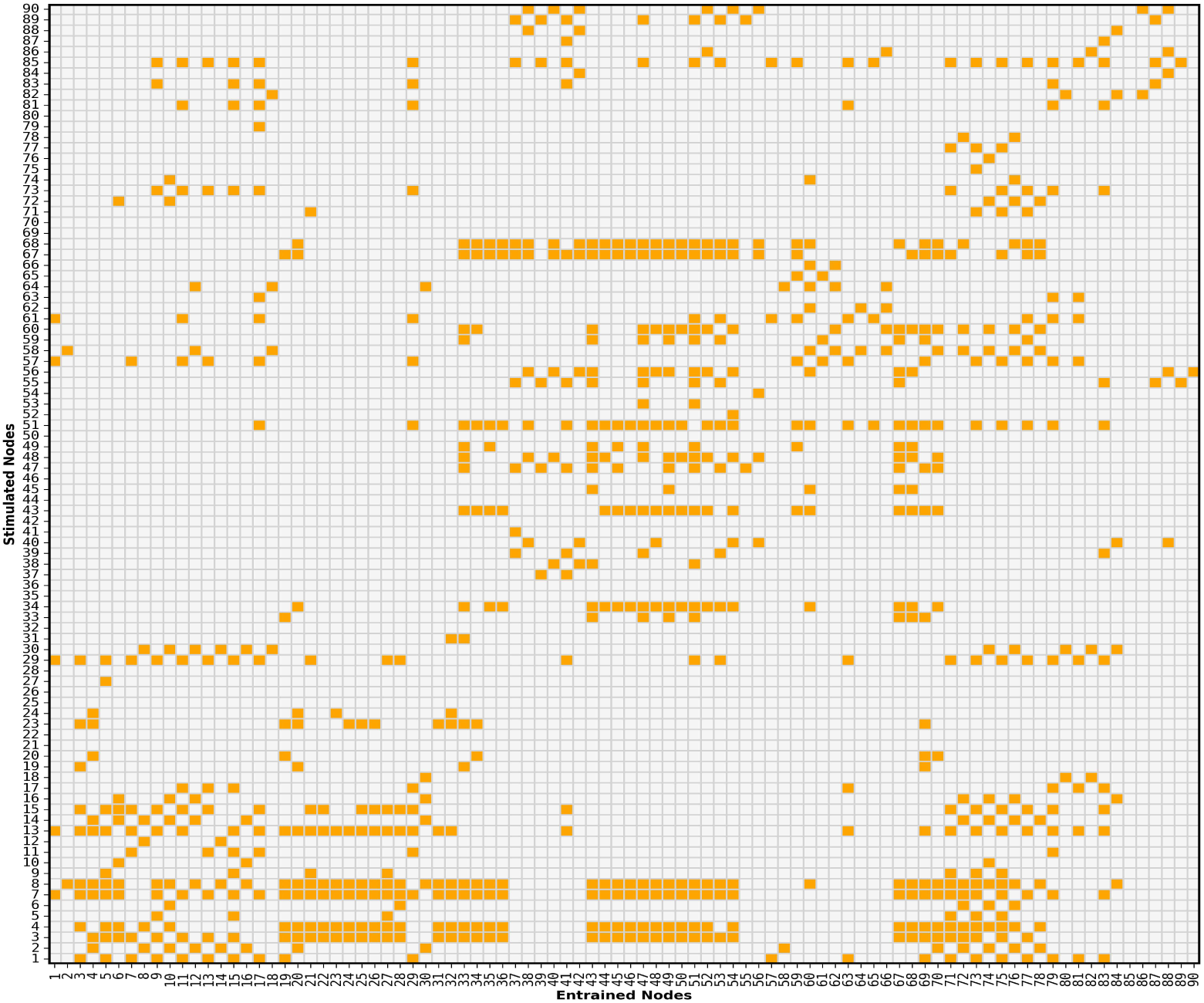
Entrainment of the nodes by stimulating each node of the network at the point of maximum entropy (K=4, MD=21) and for a frequency of 13 Hz. To check the correspondence between the numbers and the AAL90 regions see S5 Table

**S7 Fig.**
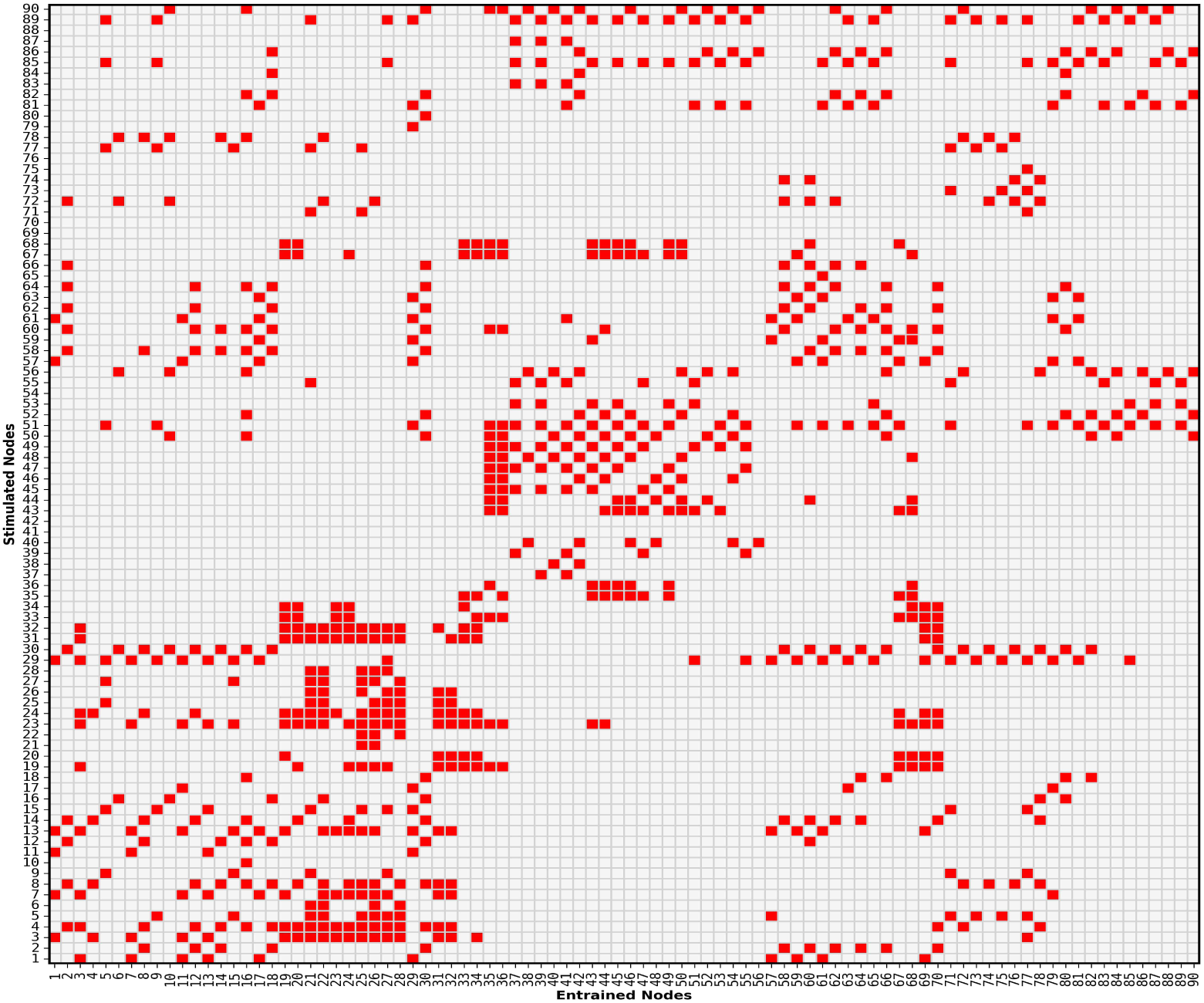
Entrainment of the nodes by stimulating each node of the network at the point of maximum entropy (K=4, MD=21) and for a frequency of 29.4 Hz. To check the correspondence between the numbers and the AAL90 regions see S5 Table

**S8 Fig.**
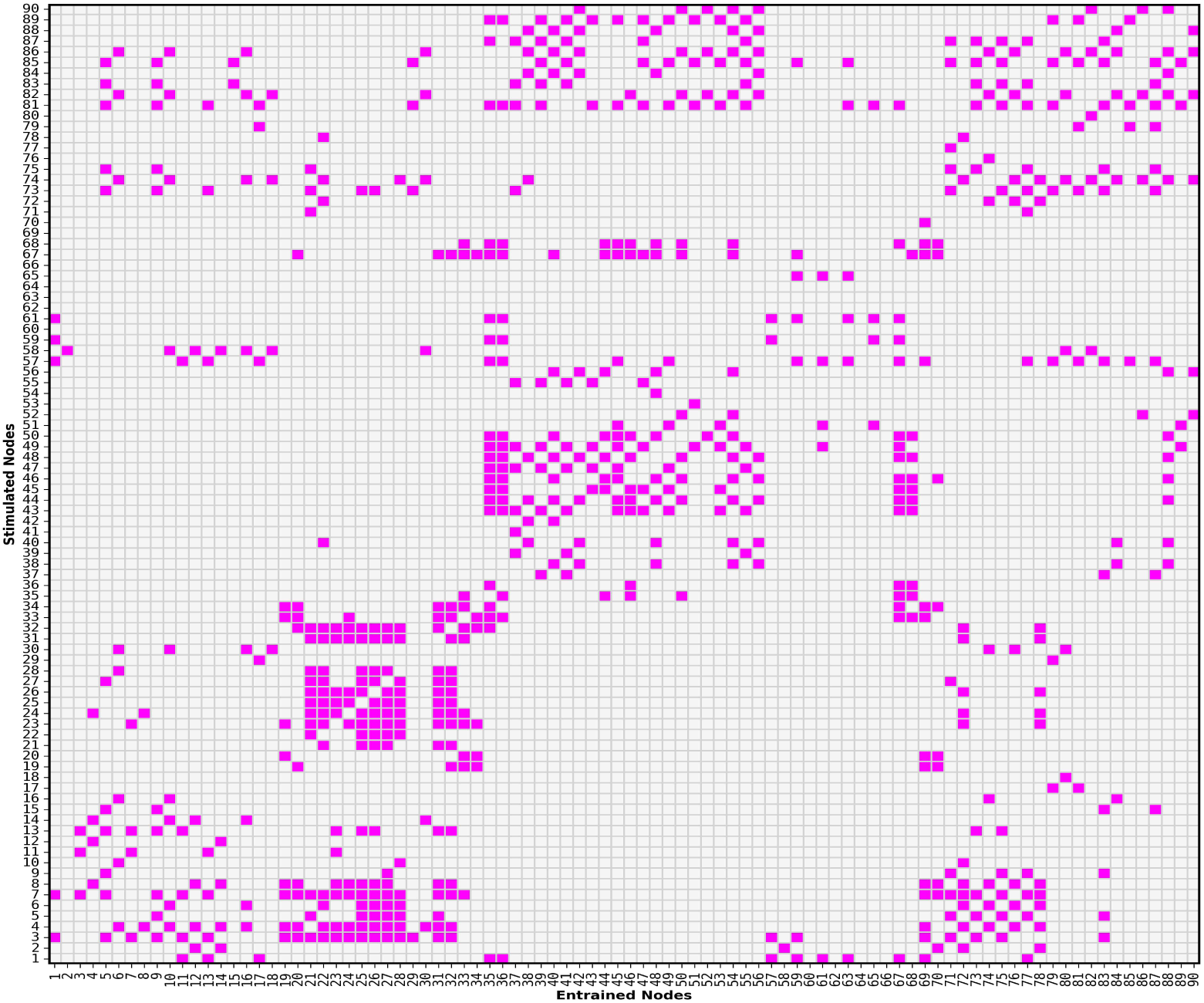
Entrainment of the nodes by stimulating each node of the network at the point of maximum entropy (K=4, MD=21) and for a frequency of 43 Hz. To check the correspondence between the numbers and the AAL90 regions see S5 Table

**S9 Fig.**
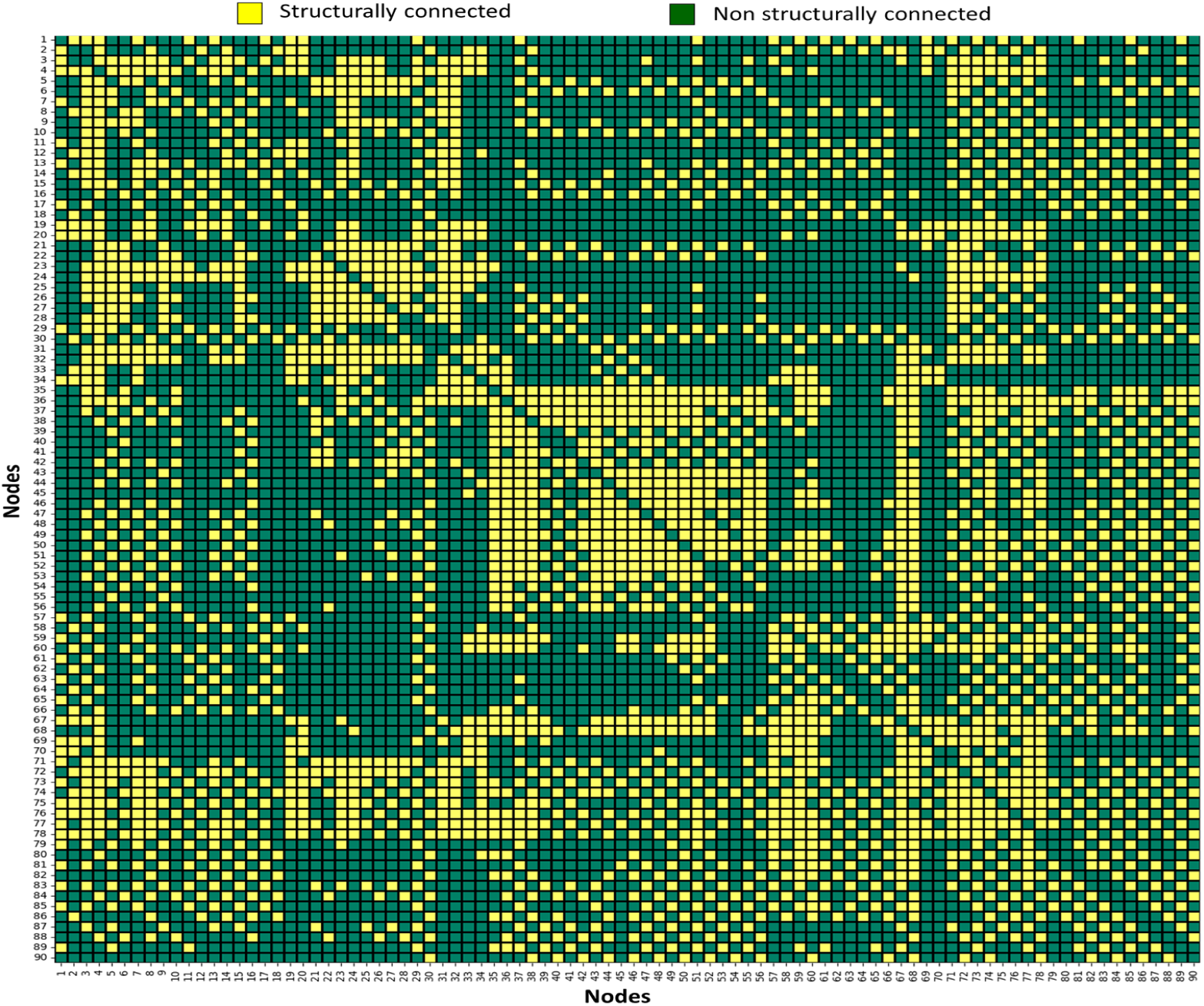
Adjacency matrix showing the structural connectivity of the 90 regions of the AAL90, with yellow indicating connected regions and green indicating unconnected regions.

**S10 Fig.**
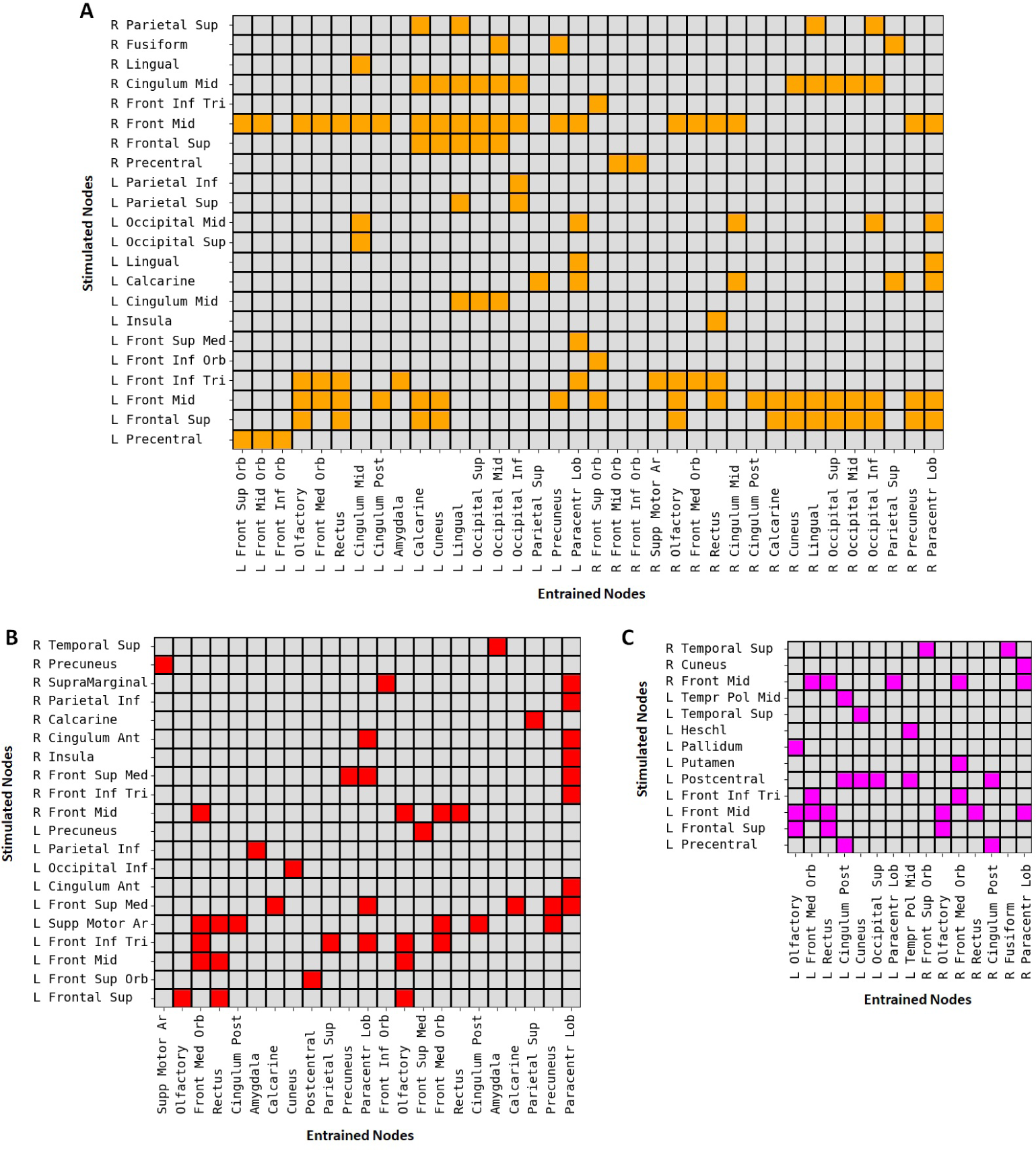
Frequency-dependent entrainment. Each panel shows the relationship between stimulated nodes (Y-axis) and indirectly entrained nodes (X-axis) at 13 Hz (A), 29.4 Hz (B), and 43 Hz (C), using the parameter set corresponding to maximum metastability ({*K*, MD} = 4, 21 ms). Entrained nodes are not directly connected to the stimulated regions but synchronize at the driving frequency. Stimulation at 13 Hz resulted in broader network propagation, while 29.4 Hz and 43 Hz induced more localized entrainment patterns.

**S11 Fig.**
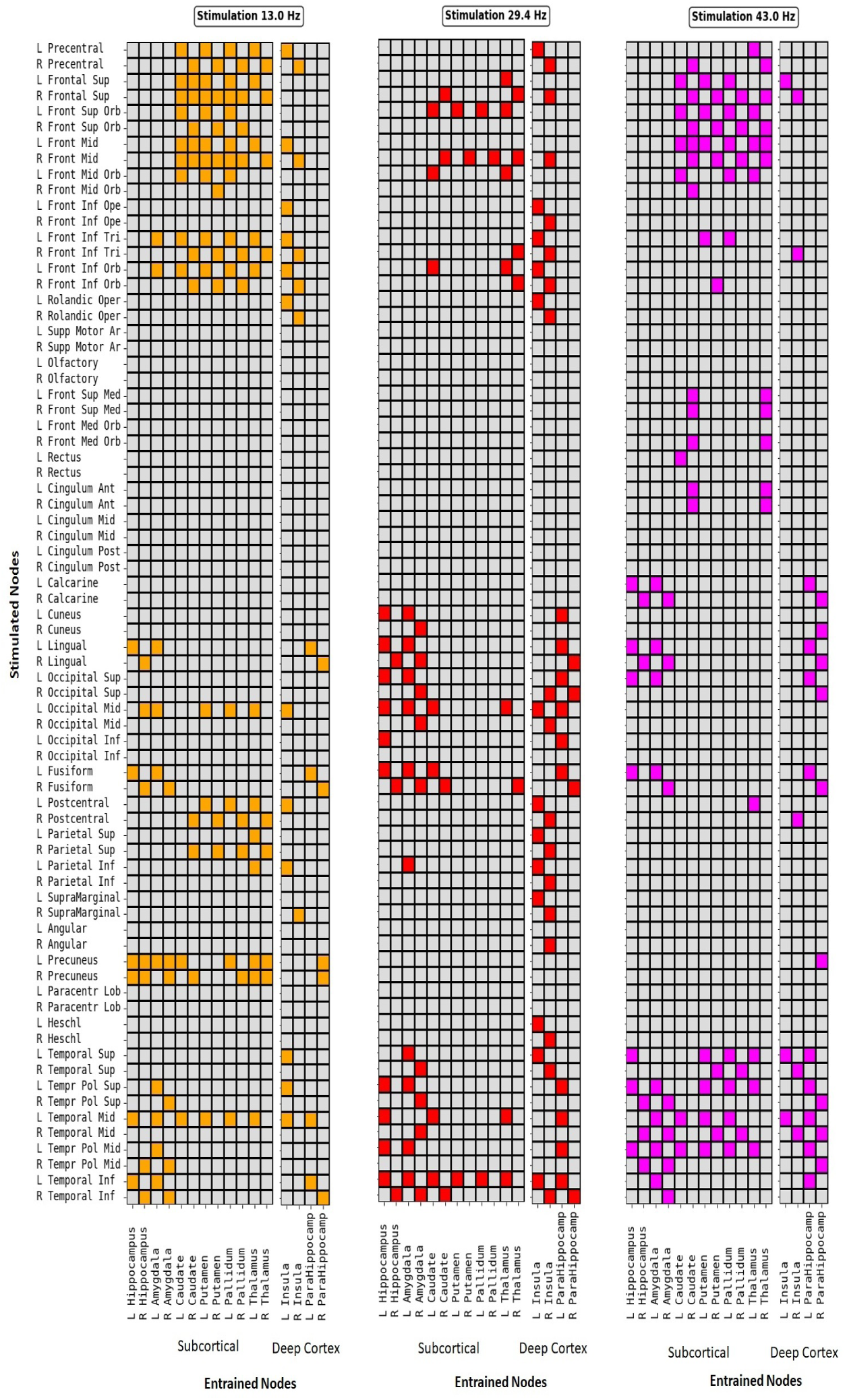
Nodes on Y-axis are stimulated at 13Hz (A), 29.4Hz (B), or 43Hz (D) at maximum metastability K,MD=4, 21ms. Nodes on the X-axis are subcortical and deep cortical structures entrained to the driving frequency. At a frequency of 13Hz entrainment reaches a widespread network, whereas 43Hz led to localized effects.

**S12 Fig.**
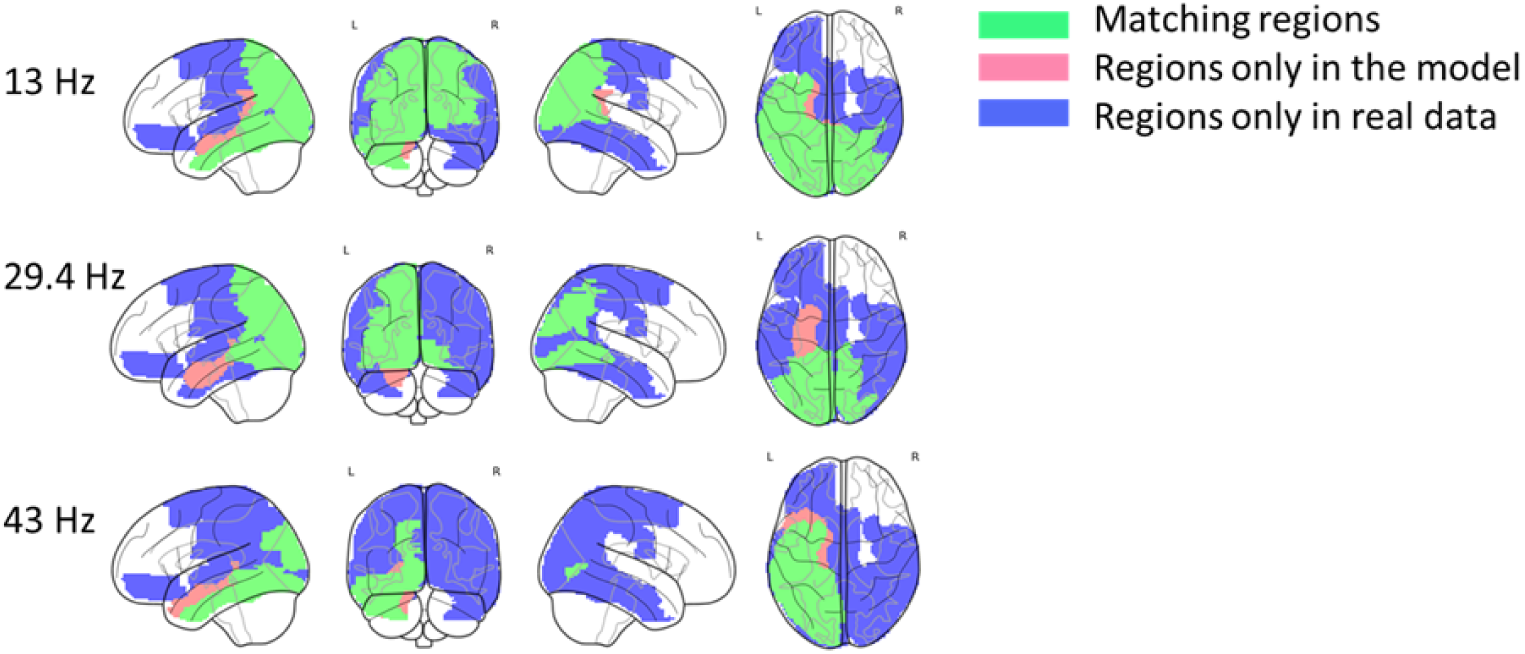
Validation of visual stimulation through spatial correspondence between empirical EEG data and model simulations across frequencies (13 Hz, 29.4 Hz, and 43 Hz). The empirical data correspond to bilateral visual stimulation, whereas the simulated data represent stimulation of the left Calcarine cortex (primary visual area). Overlapping regions are shown in green, regions exclusive to the model in red, and regions exclusive to the empirical EEG data in blue. This comparison reveals a stronger correspondence at 13 Hz — the frequency closest to the visual cortex’s preferred 10 Hz rhythm — indicating that the model successfully reproduces the frequency-dependent entrainment patterns observed in real EEG recordings .

**S13 Fig.**
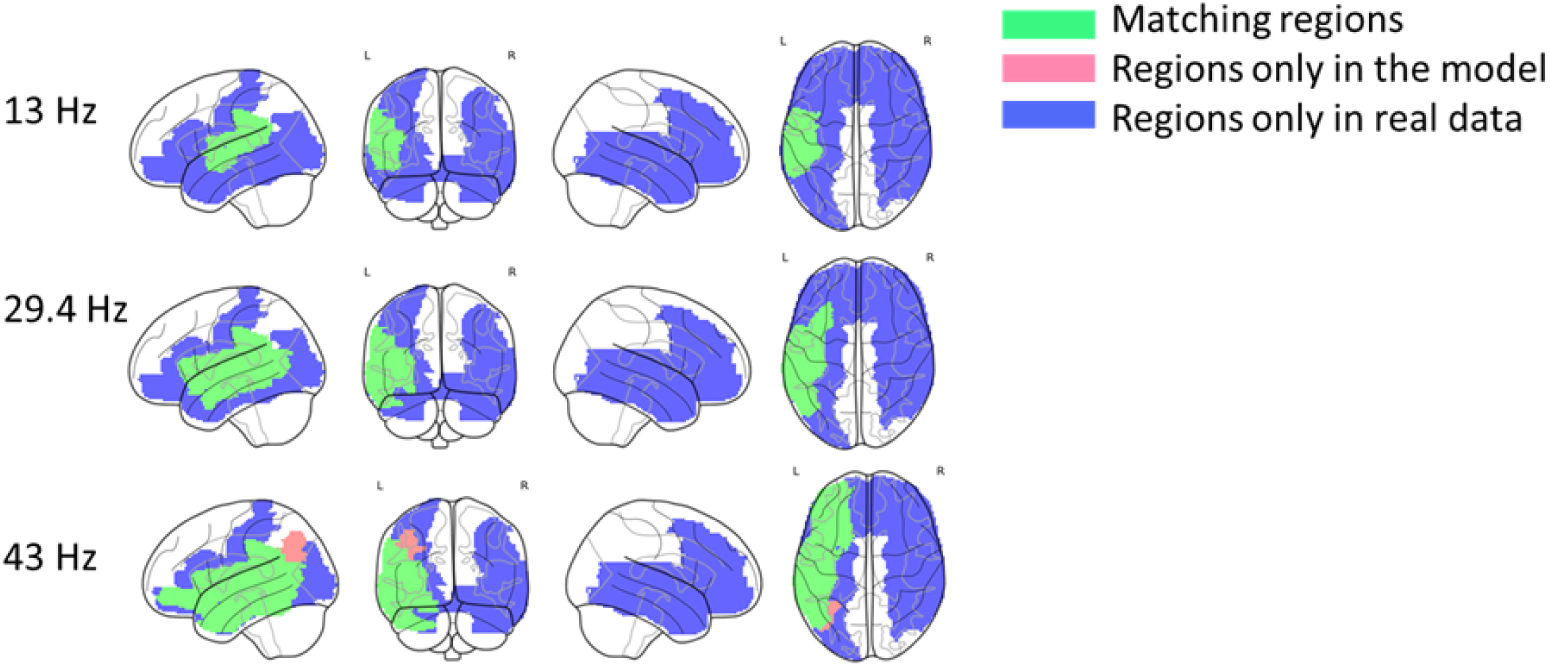
Validation of auditory stimulation through spatial correspondence between empirical EEG data and model simulations across frequencies (13 Hz, 29.4 Hz, and 43 Hz). The empirical data correspond to bilateral auditory stimulation, whereas the simulated data represent stimulation of Heschl’s gyrus (primary auditory cortex). Overlapping regions are shown in green, regions exclusive to the model in red, and regions exclusive to the empirical EEG data in blue. This comparison reveals a stronger correspondence at 43 Hz — the frequency closest to the auditory cortex’s preferred rhythm — indicating that the model accurately captures frequency-specific entrainment patterns observed in real EEG recordings. .

**S14 Table.** Validation metrics for visual stimulation. Table present the number of overlapping regions and the quantitative similarity measures between simulated and empirical data across all tested frequencies. The results show that the highest correspondence between model and experimental data occurs at 13 Hz, consistent with the frequency-selective entrainment observed in visual cortex.

| Metrics/frequencies | 13 | 29.4 | 43 |
| --- | --- | --- | --- |
| Number of matching regions | 16 | 8 | 5 |
| Pearson correlation | 0.40 ($p_{value} = 1.47 \times 10^{-4}$) | 0.30 ($p_{value} = 5.59 \times 10^{-3}$) | 0.38 ($p_{value} = 3.86 \times 10^{-4}$) |
| Cosine similarity | 0.50 | 0.33 | 0.44 |

**S15 Table.** Validation metrics for auditory stimulation. Table present the number of overlapping regions and the quantitative similarity measures between simulated and empirical data across all tested frequencies. The results show that the highest correspondence between model and experimental data occurs at 43 Hz, consistent with the frequency-selective entrainment observed in auditory cortex.

| Metrics/frequencies | 13 | 29.4 | 43 |
| --- | --- | --- | --- |
| Number of matching regions | 3 | 5 | 9 |
| Pearson correlation | 0.17 ($p_{value} = 1.12 \times 10^{-1}$) | 0.22($p_{value} = 4.14 \times 10^{-2}$) | 0.37 ($p_{value} = 637 \times 10^{-4}$) |
| Cosine similarity | 0.25 | 0.30 | 0.43 |

## Author Contributions

**Conceptualization:** Mónica Otero, Wael El-Deredy.

**Data curation:** Mónica Otero, Elida Poo, Felipe Torres, Caroline Lea-Carnall, Alejandro Weinstein.

**Formal analysis:** Mónica Otero, Elida Poo, Felipe Torres, Cristobal Mendoza, Caroline Lea-Carnall, Alejandro Weinstein, Pamela Guevara, Pavel Prado, Joana Cabral.

**Funding acquisition:** Wael El-Deredy.

**Investigation:** Mónica Otero, Elida Poo, Felipe Torres, Cristobal Mendoza, Caroline Lea-Carnall, Alejandro Weinstein, Pamela Guevara, Pavel Prado.

**Methodology:** Mónica Otero, Elida Poo, Felipe Torres, Cristobal Mendoza, Caroline Lea-Carnall, Alejandro Weinstein, Pamela Guevara, Jesús Cortés, Pavel Prado, Joana Cabral.

**Project administration:** Mónica Otero, Wael El-Deredy.

**Resources:** Mónica Otero, Wael El-Deredy, Pamela Guevara.

**Software:** Elida Poo, Felipe Torres, Cristobal Mendoza, Caroline Lea-Carnall.

**Supervision:** Mónica Otero, Wael El-Deredy.

**Validation:** Mónica Otero, Elida Poo, Felipe Torres, Caroline Lea-Carnall, Alejandro Weinstein, Pamela Guevara, Jesús Cortés, Pavel Prado, Joana Cabral.

**Visualization:** Elida Poo, Felipe Torres, Cristobal Mendoza, Caroline Lea-Carnall, Alejandro Weinstein.

**Writing – original draft:** Mónica Otero, Elida Poo

**Writing – review & editing:** Mónica Otero, Elida Poo, Felipe Torres, Cristobal Mendoza, Caroline Lea-Carnall, Alejandro Weinstein, Pamela Guevara, Jesús Cortés, Pavel Prado, Joana Cabral, Wael El-Deredy.

## Notes

### Competing Interest Statement

The authors have declared no competing interest.

### Summary of Updates

This revised version includes updates to the validation section, updated supplementary files, and a full review of the manuscript to improve clarity

